# Identification of State Markers in Anorexia Nervosa: Replication and Extension of Inflammation Associated Biomarkers Using Multiplex Profiling in Anorexia Nervosa and Atypical Anorexia Nervosa

**DOI:** 10.1101/2023.06.30.547289

**Authors:** Lauren Breithaupt, Laura M. Holsen, Chunni Ji, Jie Hu, Felicia Petterway, Megan Rosa-Caldwell, Ida A.K. Nilsson, Jennifer J. Thomas, Kyle A. Williams, Regine Boutin, Meghan Slattery, Cynthia M. Bulik, Steven E. Arnold, Elizabeth A. Lawson, Madhusmita Misra, Kamryn T. Eddy

## Abstract

Proteomics provides an opportunity for detection and monitoring of anorexia nervosa (AN) and its related variant, atypical-AN (atyp-AN). However, research to date has been limited by the small number of proteins explored, exclusive focus on adults with AN, and lack of replication across studies. This study performed Olink Proseek Multiplex profiling of 92 proteins involved in inflammation among females with AN and atyp-AN (N = 64), all ≤ 90% of expected body weight, and age-matched healthy controls (HC; N=44). After correction for multiple testing, nine proteins differed significantly in the AN/atyp-AN group relative to HC group (*lower* levels: CXCL1, HGF, IL-18R1, TNFSF14, TRANCE; *higher* levels: CCL23, Flt3L, LIF-R, MMP-1). The expression levels of three proteins (*lower* IL-18R1, TRANCE; *higher* LIF-R) were uniquely disrupted in females with AN. No unique expression levels emerged for atyp-AN. Across the whole sample, twenty-one proteins correlated positively with BMI (ADA, AXIN1, CD5, CD244, CD40, CD6, CXCL1, FGF-21, HGF, IL-10RB, IL-12B, IL18, IL-18R1, IL6, LAP TGF-beta-1, SIRT2, STAMBP, TNFRSF9, TNFSF14, TRAIL, TRANCE) and six (CCL11, CCL23, FGF-19, IL8, LIF-R, OPG) were negatively correlated with BMI. Overall, our results replicate the prior study demonstrating a dysregulated inflammatory status in AN, and extend these results to atyp-AN (AN/atyp-AN all ≤ 90% of expected body weight). Of the 27 proteins correlated with BMI, 18 were replicated from a prior study using similar methods, highlighting the promise of inflammatory protein expression levels as biomarkers of disease monitoring. Additional studies of individuals across the entire weight spectrum are needed to understand the role of inflammation in atyp-AN.

---

Anorexia nervosa (AN) and its related variant (i.e., atypical anorexia nervosa; atyp-AN) are among the deadliest psychiatric disorders. The primary distinction between AN and atyp-AN is that AN is marked by significant low weight (BMI<18.5), while those with atyp-AN maintain a weight within or above normal limits.^1^ Although significant low-weight is absent in atyp-AN, psychiatric and medical severity are comparable.^2, 3^ In fact, people with atyp-AN comprise at least one third of those with eating disorders receiving inpatient level of care due to medical instability.^3, 4^ The hallmark symptom shared across AN and atyp-AN—relentless caloric restriction—drives the associated morbidity and high rates of hospitalization. Although eventual recovery from these illnesses may occur,^5^ full recovery becomes much less likely as the illnesses progress,^6, 7^ underscoring the need for early intervention. However, AN and atyp-AN often go undetected by providers and family members for several years,^8^ and due to the egosyntonic nature of the illnesses, individuals rarely initiate treatment on their own.^9^ Diagnosis can be challenging as it rests on measured body weight (for which diagnostic weight thresholds are imperfectly defined) and self-reported symptoms (which those with AN/atyp-AN frequently minimize or fail to recognize). Particularly when validated and measured among individuals in the adolescent age range, corresponding to the initial emergence of AN/atyp-AN symptomatology,^10, 11^ biomarkers could aid in illness detection and differential diagnosis, inform our mechanistic understanding of these illnesses, and guide the conceptualization of outcomes. Thus far, studies on protein biomarkers in AN have all focused on adults with AN, usually years after the disorder emerged and evolved.

Inflammation is a regulatory process that occurs in response to injury, stress, or infection, and is widely implicated across psychiatric disorders.^12–15^ Inflammation may result from either physiological or psychological stressors and is influenced by the production of pro– and anti-inflammatory chemical messengers or networks. The role of inflammation in AN/atyp-AN etiology and symptom maintenance is complex. Adipocytes, the cell responsible for the storage of fat, are capable of producing inflammatory messengers and cytokines (i.e., tumor necrosis factor α [TNFα], interleukin [IL] 1 and 6), and changes in fat composition during states of obesity or cachexia alter expression of these molecules.^16–20^ Cytokines also play a role in appetite regulation and eating behavior,^21–23^ which may contribute to the reluctance to undergo treatment in AN and serve as a state marker of low-weight that maintains AN. Results from meta-analyses and narrative review suggest AN displays a unique profile of inflammatory molecule expression compared to primary malnutrition, with increases in circulating TNFα, IL1ß, IL6, and TNF-receptor-II and decreases C-reactive protein (CRP) and IL6-receptor expression.^24–26^ As such, inflammation protein expression levels may serve as a state-specific biomarker of AN, associated with the low-weight status.

A recent study comparing the largest battery of plasma inflammatory markers in women with active AN (N = 113), recovered from AN (N = 113), and healthy normal-weight controls (N = 114) highlighted the promise of inflammation protein expression levels as state-specific makers of AN, associated only with low-weight status. In this study, investigators observed a markedly different proteomic plasma profile of inflammatory markers in women with AN compared to both controls and women who had recovered from AN, whereas there were no differences between those who had recovered and healthy controls (HCs). The proteins identified were also correlated with BMI, suggesting that aberrant inflammatory profile is a state marker associated with AN and/or low BMI. Notably, because individuals with atyp-AN were not included in the study, parsing whether those inflammation proteins were unique to AN and/or independent of low-weight status (i.e., present in both AN and atyp-AN, but not in either group when recovered) was not possible.

To determine whether inflammatory protein markers represent useful clinical biomarkers, several follow-up steps are necessary. First, examination of inflammatory profiles in those with AN and those with atyp-AN compared to HCs is needed. This work could inform whether inflammatory markers reflect restrictive eating disorders more broadly (i.e., AN and atyp-AN), distinguishing them from HCs, or whether specific biomarkers could aid in differential diagnosis between AN and atyp-AN. Further, understanding whether and when inflammation normalizes would inform treatment targets and staging for both AN and atyp-AN. Finally, AN and atyp-AN most commonly occur in adolescents and young adults,^10, 11^ and the prior study sample comprised adults with AN. As inflammation proteins differ significantly across age ranges,^27^ studying adolescents with AN and atyp-AN is a priority.

The current study quantified inflammatory proteins in a mixed sample of female adolescents with AN, atyp-AN, and age-matched healthy controls. We included individuals with restrictive eating disorders (AN/atyp-AN) who had ≤ 90% of expected body weight. This study was designed to replicate and expand upon the findings of Nilsson et al., 2020, hypothesizing that (1) The combined AN/atyp-AN groups will display a unique profile of inflammatory protein expression compared to healthy controls. Next, to understand the contribution of low-weight status to the inflammatory protein expression, we hypothesized that (2) certain markers of immune dysregulation would be unique to AN (evident in comparisons of AN vs. HC and not in atyp-AN vs. HC). Across the entire group, we expected to see (3) strong relationships between immune proteins and BMI. To replicate findings from Nilsson, we utilized the same multiplex inflammation panel, Olink Proteomics, to measure concentrations of 92 preselected inflammation-related proteins. To test the robustness of our findings, we examined the impact of covariates, such as smoking,^28, 29^ antihistamine use,^30^ antidepressant use,^31^ and depression/anxiety comorbidity,^15^ which are known to be associated with inflammation, but that have not been previously considered.^32^

## 2. Material and methods

### 2.1 Sample

Data were derived from two parent studies described in ^33, 34^ that assessed food motivation pathways as mediators of eating disorder trajectories. The sample included females diagnosed with AN and atyp-AN, who were all ≤ 90% of median BMI (based on the 50th percentile of body mass index [BMI: weight (kg)/height(m)^2^] for age and gender) and HC. Participants with AN/atyp-AN and HC were enrolled in the study between 4/1/2014 and 1/18/2021 at Massachusetts General Hospital (MGH), Boston, USA with Institutional Review Board approval. All participants provided written informed consent or parental consent with assent from minors younger than 18 years. Participants were recruited from clinical programs at MGH and beyond, including the MGH Eating Disorders Clinical and Research Program, specialized treatment centers in the greater Boston area, student health centers at local universities, and community advertisements. In this report, we included females with AN, atyp-AN, and HC who were enrolled in the parent studies and had stored plasma samples at the baseline visit (N = 108). Participant criteria and study procedures are included in **<u>eMethods Supplement</u>** and reported elsewhere.^33^

### 2.2 Participants

We examined females aged 10 to 22 years with *Diagnostic and Statistical Manual of Mental Disorders* (Fifth Edition) (*DSM-5*) AN or atyp-AN, or HCs with no lifetime history of any psychiatric diagnoses or eating disorders, and with similar pubertal (Tanner) stages. The DSM-5 defines atyp-AN as meeting all criteria for AN, except for the low-weight criterion (operationalized in this study as a body mass index (BMI) less than 18.5 for adults, or a BMI-for– age percentile of at or below 10.99 for those under 18 (Center for Disease Control and Prevention [CDC]). Screening for AN/atyp-AN included clinical evaluation to confirm the lack of any organic condition that could account for the low body weight. Study participants had no history of diabetes mellitus, gastrointestinal tract surgery, recent systemic hormone use, or pregnancy, which could impact levels of inflammatory proteins. No participants had taken non-steroidal anti-inflammatory drugs (NSAIDs) on the day of the collection, and we recorded NSAIDs use over the past 2 months, which was then used as a covariate in secondary models (see 2.4 Statistical Analysis Plan). Participants were medically and psychiatrically stable with sufficiently mitigated imminent risk of harm, without dangerous and active medical issues (e.g., severe or rapidly progressing electrolyte imbalances or anemia) or acutely elevated psychiatric risk (e.g., active suicidality). See the eMethods Supplement for details of the selection criteria.

### 2.3 Blood sampling and processing

We collected fasting blood at around 8:45 AM (at least 8 hours of fasting with the exception of water intake) by trained nursing staff using EDTA tubes. After centrifugation at 4° C, the collected plasma samples were stored at –80° C. Samples were processed by Olink’s Target Inflammation 96 panel, which utilizes proximity extension assay (PEA) technology with pairs of oligonucleotide-labeled antibodies binding to the target protein, followed by amplification using polymerase chain reaction (PCR).^35^ The panel can quantify 92 inflammation-related proteins simultaneously and reports the protein concentrations in a relative log_2_ scale called normalized protein expression (NPX), which is calculated from the cycle threshold (Ct) values, inverted, and normalized by Olink to minimize both intra– and inter-assay variation. A difference of 1 NPX unit approximates a doubling of the protein concentration. We assessed the normality of protein concentrations by visually assessing the quantile–quantile plots. Given that proteins are reported in a log_2_ scale and the majority (see Supplementary eFig. 1) of analyzed proteins were normally distributed, no additional normalization procedures were applied. In addition, Olink reports the limit of detection (LOD) for each assay estimated from negative controls (i.e., contain buffer only) plus three standard deviations. Samples that do not pass quality control are flagged with a quality control warning (QC warning). For detailed information on LOD and QC procedures, see ^35^ and (https://www.olink.com/resources-support/white-papers-from-olink/).

**Figure 1.**
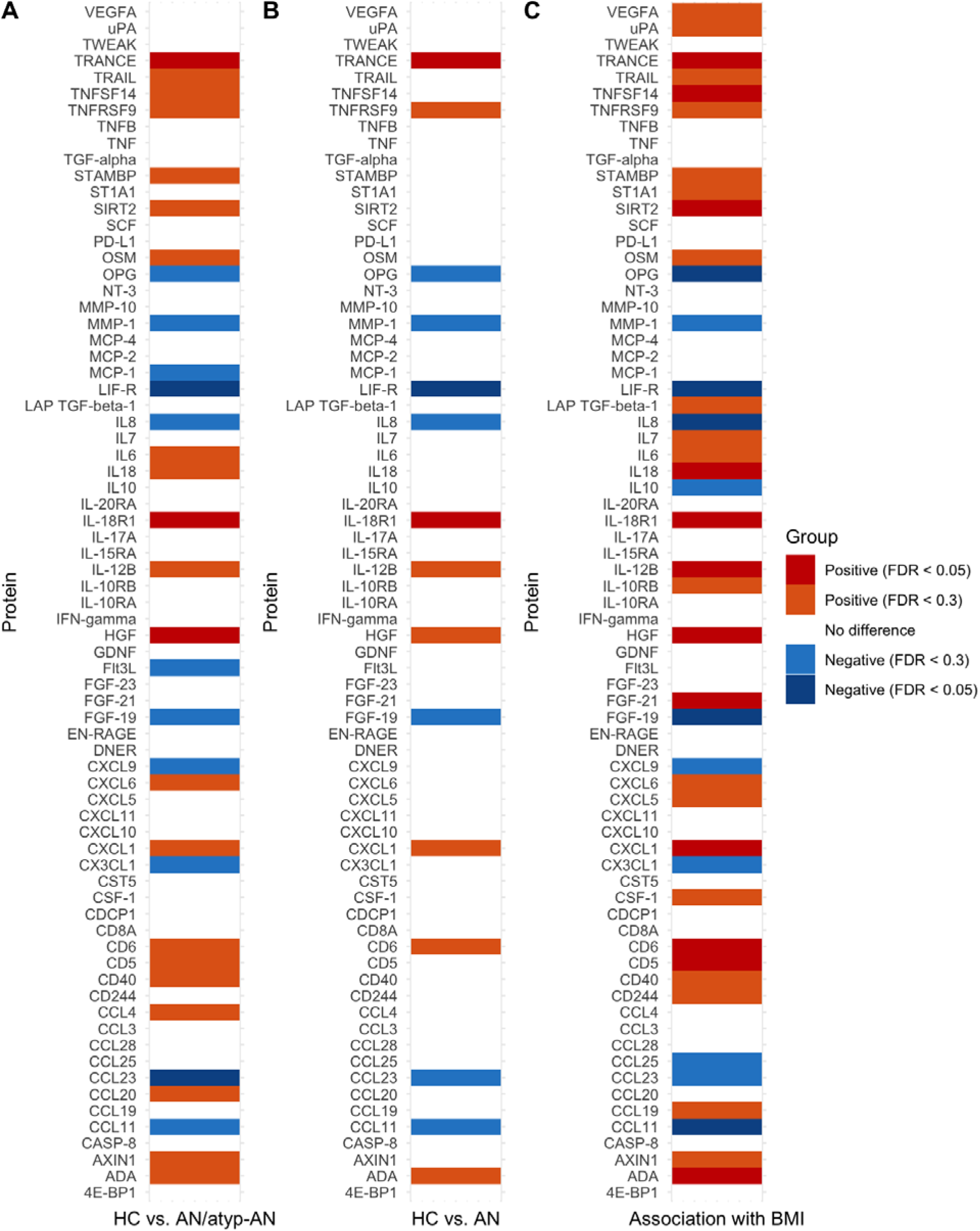
Plasma protein concentrations that differ between HC vs. AN/atyp-AN (A), differ between HC vs. AN (B), and are associated with BMI (C), after adjusting for age (Model 1). A positive result indicates either the protein has a higher plasma concentration in the HC group or is positively associated with BMI. No significant results were observed between HC vs. atyp-AN or between AN vs. atyp-AN. AN = anorexia nervosa, atyp-AN = atypical anorexia nervosa, HC = healthy control.

### 2.4 Statistical analyses

Statistical analyses were performed using R version 4.2.2 (R Core Team, 2022). All 108 samples passed the general Olink QC. Protein values lower than the LOD were replaced with the corresponding LOD value. Consistent with Nilsson et al. (2020), proteins with over 20% of data lower than the LOD were excluded from the analyses, leaving us with a total of 73 proteins for analysis. Moreover, we visually inspected the summary plot from the principal component analysis (PCA) to identify outliers.

We conducted multiple linear regression analyses to compare group differences for each protein between AN/atyp-AN vs. HC (Aim 1) and AN vs. atyp-AN vs. HC (Aim 2), followed by associations between BMI z-score and each protein across the entire group (Aim 3). We tested two primary models for each aim: in Model 1, we included only age as a covariate; in Model 2, we included the following variables as covariates: age, smoking status, body temperature, any antihistamine use, any psychiatric medication use, and any psychiatric comorbidity. We adjusted multiple comparisons using the Benjamini-Hochberg correction.^36^ An FDR <0.05 was considered statistically significant, and an FDR of 0.06-0.1 as near significance was considered marginally significant.

Following our primary analyses (Model 1, Model 2), we conducted a series of secondary multiple linear regression models to test the robustness of the primary analyses, including factors that have been shown to influence inflammation levels, i.e., smoking status, acute sickness (high body temperature), psychiatric severity, self-reported state levels of depression/anxiety, antidepressant use, or antihistamine use. All models are described in Table 1. In Secondary Analysis 1 (Model S1), we excluded four smoking subjects and accounted for age. In Secondary Analysis 2 (Model S2), we excluded eight participants with a body temperature higher than 99° F and accounted for age. Secondary Analysis 3 (Model S3) controlled for age and the number of psychiatric diagnoses. Given that depression and anxiety were highly correlated, Secondary Analysis 4 (Model S4) accounted for age and only the anxiety T-score measured by the State-Trait Anxiety Inventory (STAI)^37^ (Pearson correlation coefficient between anxiety t-score and depression t-score: r = 0.81, p < 0.001). Secondary Analysis 5 (Model S5) accounted for age and antidepressant use, whereas Secondary Analysis 6 (Model S6) accounted for age and NSAIDs/antihistamine use over the past 2 months. All R code can be downloaded from the projects Github page <u>here</u>.

**Table 1.** Model Details.

| Model Details |  |  |
| --- | --- | --- |
|  | Covariate(s) | Subjects Removed |
| **Main Models:** |  |  |
| **Model 1** | Age |  |
| **Model 2** | Age, smoking status, body temperature, any antihistamine use, any psychiatric medication use, and any psychiatric comorbidity |  |
| **Secondary Models:** |  |  |
| **S1** | Age | Smoking (n = 4) |
| **S2** | Age | Body temperature greater than 99° F (n = 8) |
| **S3** | Age, psychiatric comorbidity |  |
| **S4** | Age, anxiety/depressive symptoms |  |
| **S5** | Age, antidepressant use |  |
| **S6** | Age, antihistamine use |  |

## 3. Results

We included a total of 108 females (17 [16%] Asian, 89 [82%] White, 2 [2%] Other; 5 [5%] Hispanic), of whom 38 had AN (median [IQR] age, 20.0 [18.5 – 21.1] years), 26 had atyp-AN (median [IQR] age, 19.0 [15.8 – 21.1] years), and 44 were HCs (median [IQR] age, 18.8 [15.6 – 20.3] years), with similar percentages of premenarchal participants in each group. As expected, BMI z score significantly differed by group (BMI z score mean (SD): AN = –2.26 (0.741), atyp-AN = –0.864 (0.264), HC = 0.118 (0.556); F(2, 105) = 171.1, p < 0.001). Individuals with AN were significantly older than HC (Tukey’s HSD test, p = 0.01); however, there were no significant age differences between AN and atyp-AN or atyp-AN and HC. Importantly, AN and atyp-AN did not differ in duration of illness, frequency of binge/purge symptoms, number of psychiatric comorbidities, levels of depression, levels of anxiety, or psychiatric medication use. Clinical characteristics of the sample are summarized in Table 2, and Supplementary Table1 provides more information on specific psychiatric medication use and psychiatric comorbidity. We excluded nineteen proteins (MCP-3, IL-17C, IL-2RB, IL-1 alpha, IL2, TSLP, SLAMF1, FGF-5, IL-22 RA1, Beta-NGF, IL-24, IL13, ARTN, IL-20, IL33, IL4, LIF, NRTN, IL5) from further analysis due to over 20% of invalid data (values lower than the LOD). We identified and removed two outliers from the HC group through visual inspection of the PCA summary plot (see Supplementary eFig. 2) based on data from the 73 proteins. All proteins included in the analysis are in Supplementary eTable 2.

**Figure 2.**
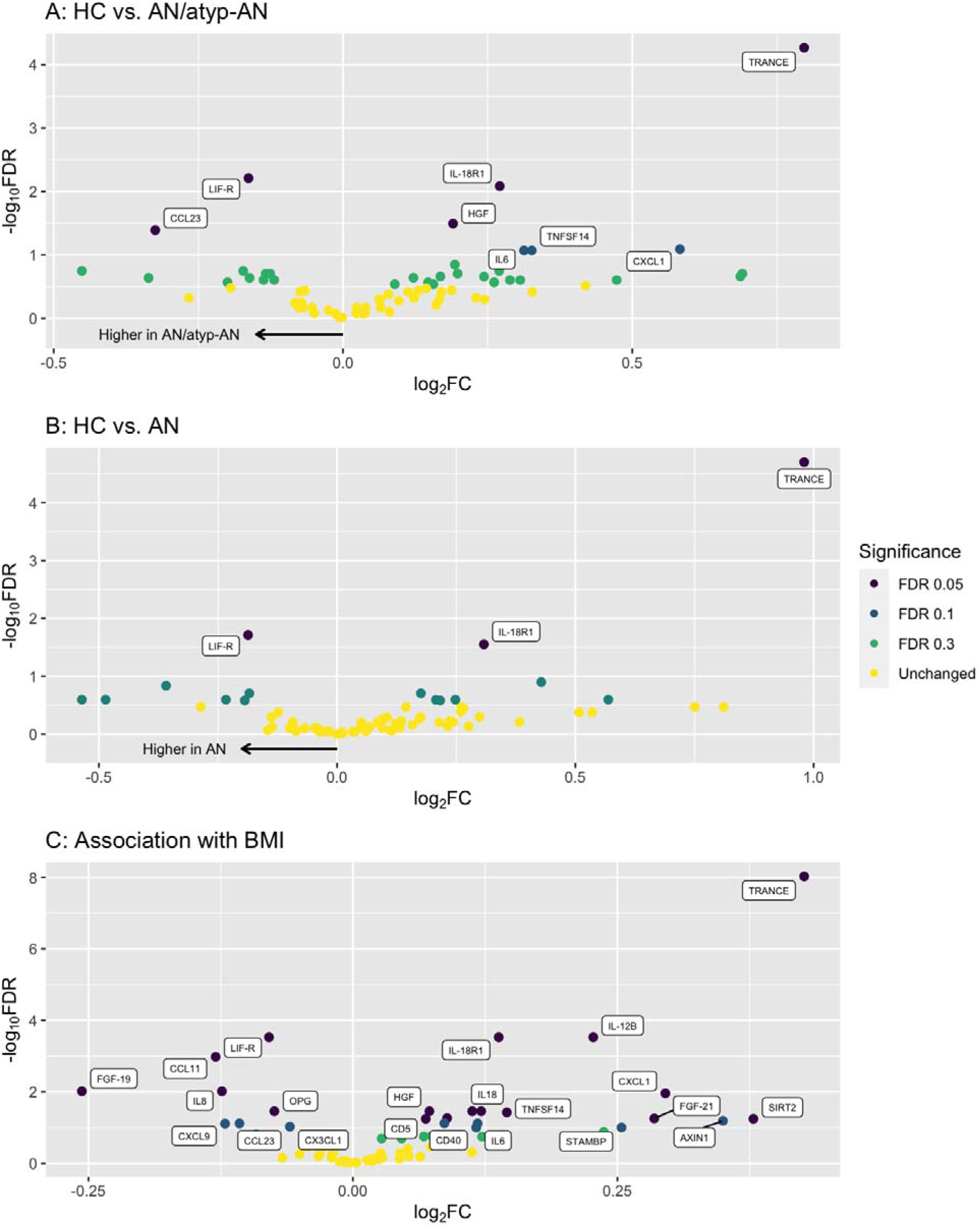
Statistical significance and log_2_ fold change (log_2_FC) for proteins that were found to be different between HC vs. AN/atyp-AN (A), different between HC vs. AN (B), and associated with BMI (C), after adjusting for age (Model 1). The x-axis represents log_2_FC. Because protein concentrations were reported in a log_2_ scale (i.e., NPX), in our analysis, log_2_FC is equivalent to the beta coefficient associated with the group (A and B), or the slope of BMI in the regression model (C). In panel C, it indicates the changes in protein concentration on a log_2_ scale for every 1 unit change in BMI. The y-axis depicts –log_10_FDR, with smaller FDR values corresponding to higher –log_10_FDR values. A value of 1.3 on the –log_10_FDR scale corresponds to an FDR of 0.05.

**Table 2.** Clinical characteristics of our sample.

|  | **AN**<br>**(N=38)** | **atyp-AN**<br>**(N=26)** | **HC**<br>**(N=44)** | **Overall**<br>**(N=108)** |
| --- | --- | --- | --- | --- |
| **Age (years)** |  |  |  |  |
| Median [IQR] | 20.0 [18.5 - 21.1] | 19.0 [15.8 - 21.1] | 18.8 [15.6 - 20.3] | 19.2 [17.3 - 20.9] |
| **Race** |  |  |  |  |
| Asian | 7 (18.4%) | 3 (11.5%) | 7 (15.9%) | 17 (15.7%) |
| White | 30 (78.9%) | 23 (88.5%) | 36 (81.8%) | 89 (82.4%) |
| **Ethnicity** |  |  |  |  |
| Hispanic | 3 (7.9%) | 0 (0%) | 2 (4.5%) | 5 (4.6%) |
| No | 35 (92.1%) | 26 (100%) | 42 (95.5%) | 103 (95.4%) |
| **BMI z-score** |  |  |  |  |
| Mean (SD) | -2.26 (0.741) | -0.864 (0.264) | 0.118 (0.556) | -0.954 (1.19) |
| **Premenarchal** |  |  |  |  |
| Yes | 2 (5.3%) | 3 (11.5%) | 6 (13.6%) | 11 (10.2%) |
| No | 36 (94.7%) | 23 (88.5%) | 38 (86.4%) | 97 (89.8%) |
| **ED Duration (years)** |  |  |  |  |
| Mean (SD) | 4.91 (3.28) | 3.67 (3.60) | NA | NA |
| Missing | 1 (2.6%) | 0 (0%) | NA | NA |
| **Binge/Purge** |  |  |  |  |
| Yes | 11 (28.9%) | 7 (26.9%) | 0 (0%) | 18 (16.7%) |
| No | 27 (71.1%) | 19 (73.1%) | 44 (100%) | 90 (83.3%) |
| **Smoking** |  |  |  |  |
| Yes | 3 (7.9%) | 1 (3.8%) | 0 (0%) | 4 (3.7%) |
| No | 35 (92.1%) | 25 (96.2%) | 44 (100%) | 104 (96.3%) |
| **Psych Meds** |  |  |  |  |
| Yes | 22 (57.9%) | 16 (61.5%) | 0 (0%) | 38 (35.2%) |
| No | 16 (42.1%) | 10 (38.5%) | 44 (100%) | 70 (64.8%) |
| **Antihistamine Use** |  |  |  |  |
| Yes | 3 (7.9%) | 5 (19.2%) | 2 (4.5%) | 10 (9.3%) |
| No | 35 (92.1%) | 21 (80.8%) | 42 (95.5%) | 98 (90.7%) |
| **Antihistamine & NSAIDs Use** |  |  |  |  |
| Yes | 6 (15.8%) | 8 (30.8%) | 9 (20.5%) | 23 (21.3%) |
| No | 32 (84.2%) | 18 (69.2%) | 35 (79.5%) | 85 (78.7%) |
| **Psych Comorbidity** |  |  |  |  |
| Yes | 23 (60.5%) | 15 (57.7%) | 0 (0.0%) | 39 (36.1%) |
| No | 15 (39.5%) | 11 (42.3%) | 44 (100.0%) | 69 (63.9%) |

AN = anorexia nervosa, atyp-AN = atypical anorexia nervosa, HC = healthy control, IQR = interquartile range, SD = standard deviation, ED = eating disorder, NSAIDs = non-steroidal anti-inflammatory drugs.

### 3.1 Inflammatory markers in combined AN/atyp-AN group vs. HC group

After adjusting for age in Model 1, we found significant differences in five proteins between the combined AN/atyp-AN group and HC group (see Fig. 1A). Specifically, relative to HC, the AN/atyp-AN group had lower plasma concentrations of TRANCE, IL-18R1, and HGF, and higher concentrations of LIF-R and CCL23. Fig. 2A illustrates the log_2_FC of protein concentrations between the two groups. After adjusting for covariates in Model 2 (age, smoking status, body temperature, antihistamine use, psychiatric medication use, and psychiatric comorbidity), we found that TRANCE, IL-18R1, and LIF-R remained significantly different between the AN/atyp-AN group and HC; additionally, MMP-1 was significantly increased in the AN/atyp-AN group.

We tested the robustness of the models through an exploratory series of complementary sensitivity analyses. We conducted six secondary analyses (Models S1-S6) to ensure that our results were not driven by potential confounding factors. In each secondary model, we adjusted for different covariates while accounting for age to ensure comparability with Model 1. TRANCE, IL-18R1, and LIF-R were consistently significant in all models (Fig. 3A). HGF and CCL23 were significant in models S1 (Smoking), S2 (Body temperature), S3 (Psychiatric comorbidity), and S6 (Antihistamine use). CXCL1 and TNFSF14 showed significance exclusively in Model S2 (Body temperature), and Flt3L showed significance only in Model S4 (Anxiety/depression severity).

**Figure 3.**
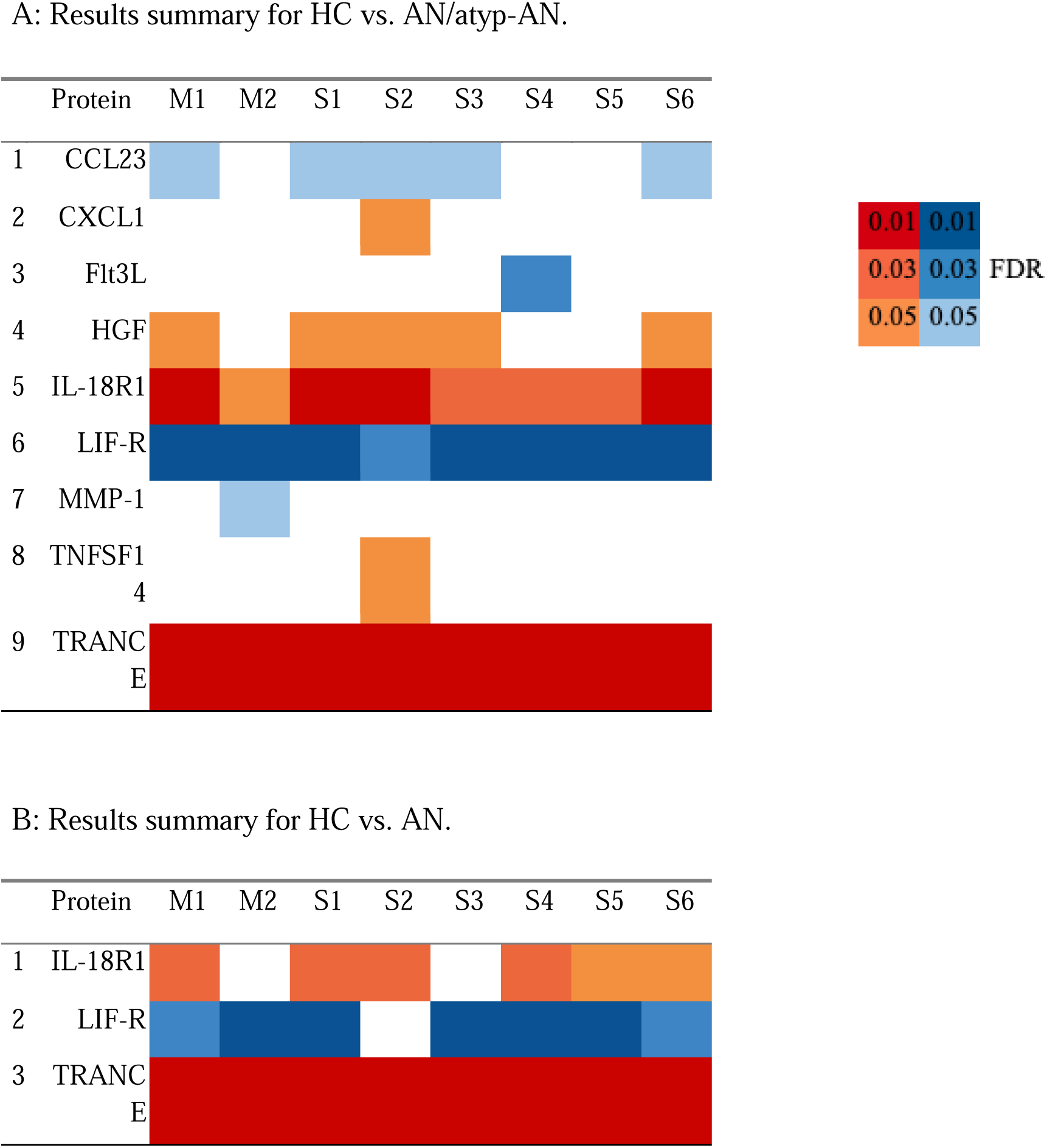
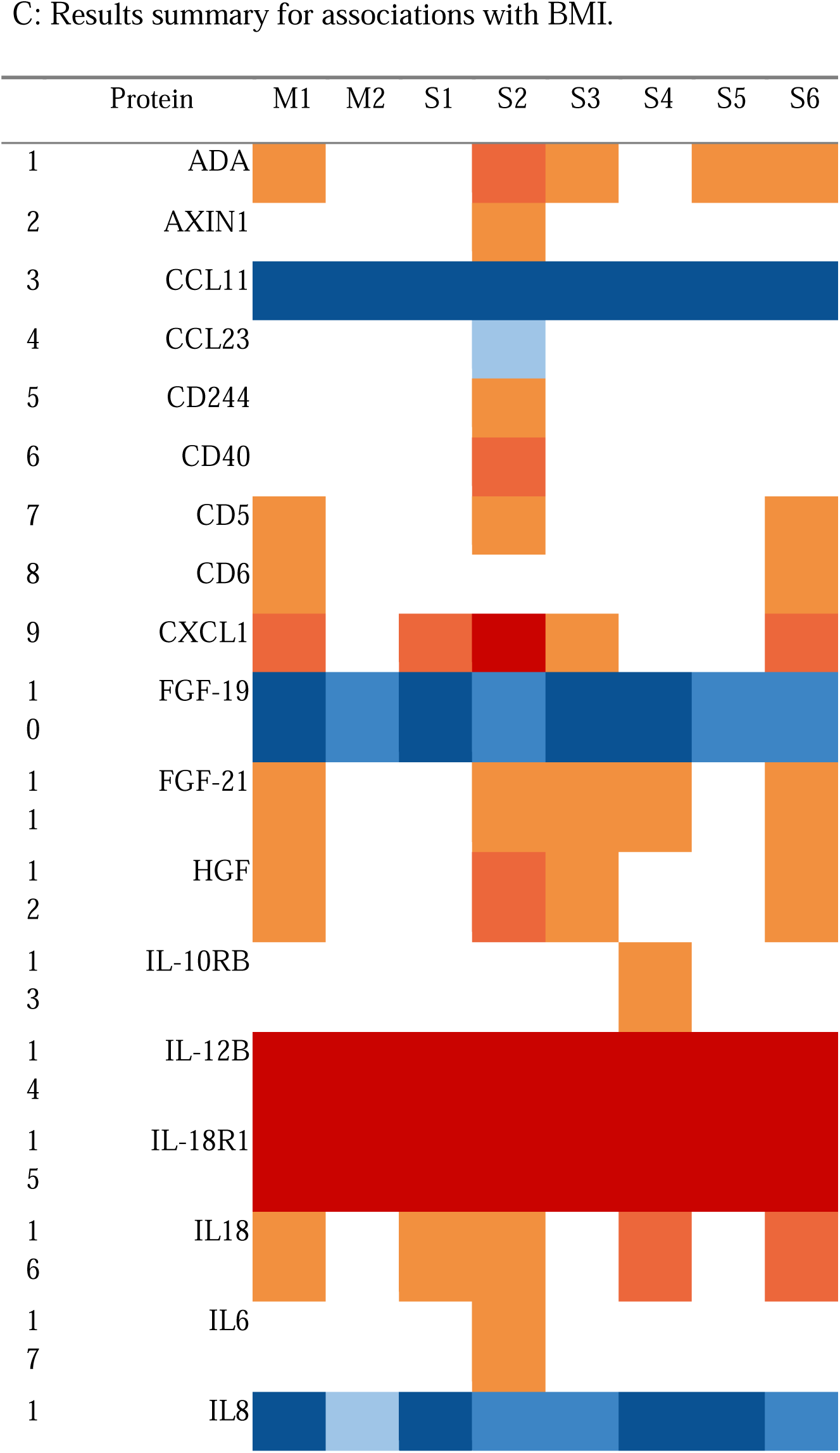
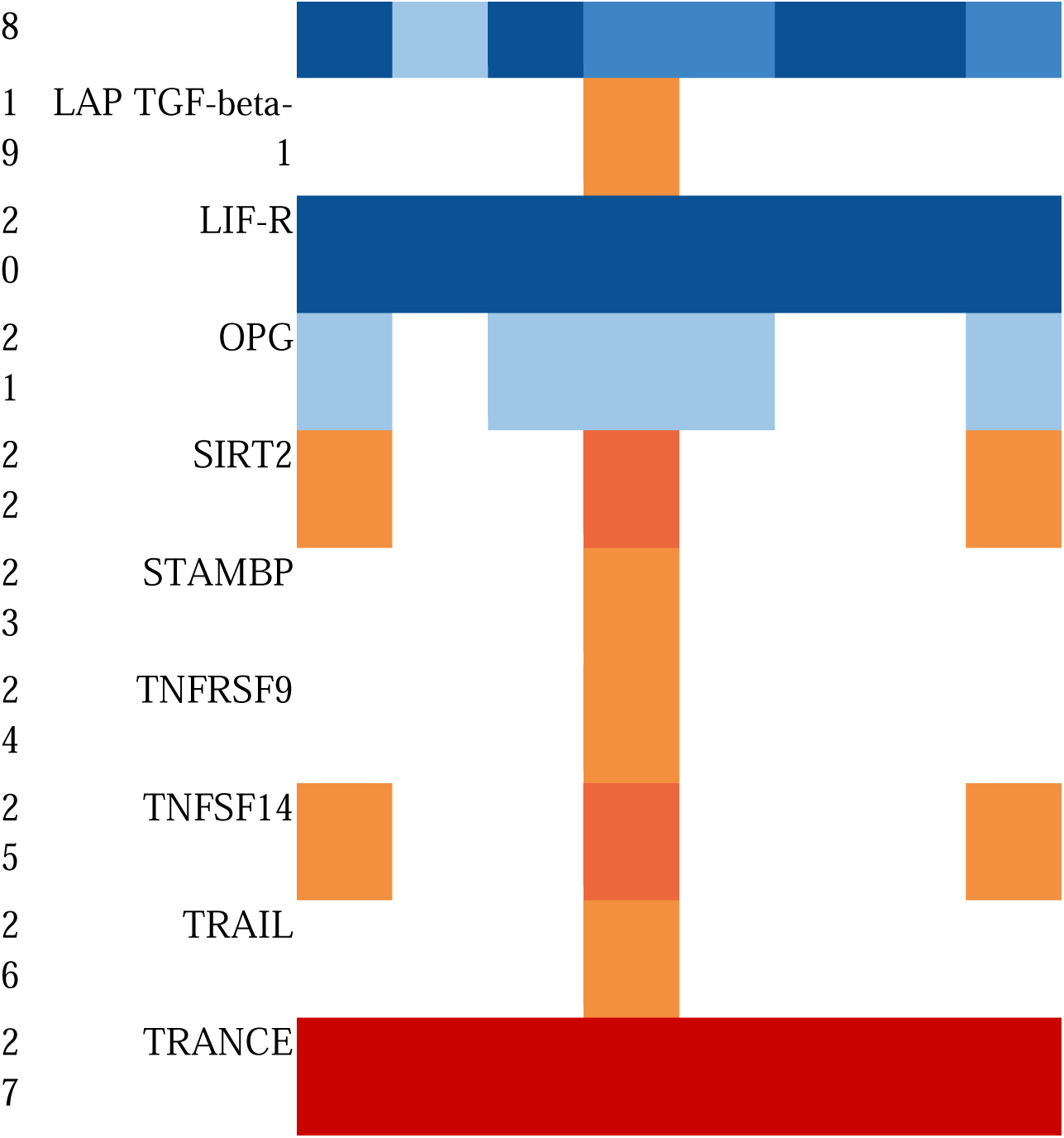
Comparison of results for HC vs. AN/atyp-AN (A), HC vs. AN (B), and association with BMI (C) across Model 1, Model 2, and Supplementary Models 1-6. Red cells represent significantly higher plasma concentrations in the HC group or significant positive associations with BMI (FDR < 0.05). Blue cells represent significantly lower plasma concentrations in the HC group or negative positive associations with BMI (FDR < 0.05). Blank cells indicate non-significance. Darker colors correspond to smaller p-values.

Across the primary and secondary models, a total of nine proteins emerged to differentiate AN/atyp-AN from HC (summarized in Fig. 3A): CCL23, CXCL1, Flt3L, HGF, IL-18R1, LIF-R, MMP-1, TNFSF14, TRANCE.

### 3.2 Inflammatory expression levels unique to low-weight AN

When comparing AN to HC in Model 1, we observed significant group differences in three proteins, specifically a lower TRANCE and IL-18R1 concentration and a higher LIF-R concentration in AN (see Fig. 1B). Fig. 2B shows the log_2_FC for protein concentrations when comparing HC vs. AN. After adjusting for covariates in Model 2 [age, smoking status, sickness (high body temperature), antihistamine use, psychiatric medication use, psychiatric comorbidity levels], TRANCE and LIF-R remained significantly different between AN and HC. Across the primary and secondary models, a total of three proteins emerged to differentiate low-weight AN from HC (summarized in Fig. 3B): TRANCE, IL18-R1, and LIF-R.

We did not observe any significant difference in protein expression when comparing AN to atyp-AN or atyp-AN to HC in our primary models. In the secondary models, levels of two proteins (LIF-R and Flt3L) were significantly higher in atyp-AN compared to HC in Model S4 where we accounted for age and anxiety t-score; no significant proteins emerged for the other secondary models.

### 3.3 Inflammatory markers and BMI across the entire sample

Across the whole sample, we analyzed the association between BMI z-score and each protein. In Model 1, we controlled for age and found that seventeen proteins were significantly associated with BMI, after correcting for multiple testing. Among them, twelve proteins (TRANCE, IL-12B, IL-18R1, CXCL1, IL18, HGF, ADA, TNFSF14, CD6, FGF-21, SIRT2, CD5) were positively associated with BMI, while five proteins (LIF-R, CCL11, FGF-19, IL8, OPG) were negatively associated with BMI. Fig. 1C shows the significance and directionality of these associations, and Fig. 2C shows the magnitude of the associations (log_2_ fold change, or beta coefficient).

In Model 2, after adjusting for age, smoking status, body temperature, antihistamine use, psychiatric medication, and psychiatric comorbidity, TRANCE, IL-12B, and IL-18R1 remained positively associated with BMI, while LIF-R, CCL11, FGF-19, and IL8 remained negatively associated with BMI. Although non-significant, ADA was positively associated with BMI at the trend level (FDR = 0.065).

We tested the robustness of the models through complementary sensitivity analyses testing secondary Models S1-S6 to ensure that our results were not driven by potential confounding factors. When testing for associations with BMI, we found that seven proteins (TRANCE, IL-12B, LIF-R, CCL11, IL8, IL-18R1, FGF-19) were significant in all models (Model 1 and S1-S6). OPG was significant in all models with the exception of Model S4 and S5 (FDR > 0.1). CXCL1 was significant in Model 1, S1, S2, S3, and S6, trend-level significant (FDR < 0.1) in Model S4, and non-significant in Model S5 (FDR > 0.1). ADA was significant in all models except for Model S1 (FDR < 0.1) and S4 (FDR > 0.1), and FGF-21 was significant in all models except for Model S1 and S5 (FDR < 0.1). IL18 was found to be significant in all models except for Model S3 (FDR < 0.1) and S5 (FDR > 0.1). In Model S2, with the exclusion of subjects with body temperatures greater than 99° F, nine other proteins (AXIN1, CCL23, CD244, CD40, IL6, LAP TGF-beta-1, STAMBP, TNFRSF9, TRAIL) were found to be significant. Model S4 accounted for anxiety severity and identified IL-10RB as significant.

Across the primary and secondary models, a total of 27 proteins were significantly correlated with BMI (summarized in Fig. 3C).

## 4. Discussion

For the first time in an adolescent sample, we analyzed 73 inflammation proteins in a cohort of females with AN, atyp-AN, and age-matched healthy controls. Consistent with our hypothesis, inflammation protein profiles were significantly different in adolescents with AN/atyp-AN compared to controls. The differences between the AN/atyp-AN and HC groups were driven by nine proteins: TRANCE, IL-18R1, HGF, LIF-R, CCL23, MMP-1, CXCL1, TNFSF14, and Flt3L. Further, we identified three proteins that uniquely distinguished low-weight AN from HC: TRANCE, LIF-R, IL-18R1. Twenty-seven proteins, including TRANCE, LIF-R, IL-18R1, HGF, CCL23, CXCL1 and TNFSF14, were correlated with BMI across the entire sample. Of note, robust associations in our extensive series of sensitivity analyses support the validity of our findings, including disrupted levels of TRANCE, IL-18R1, and LIF-R as unique to low-weight AN (level not unique in atyp-AN) and the association of these proteins with BMI. Below we discuss individual proteins identified, and their current known roles, however many of the proteins identified likely also have functions in the nervous system that may not be well studied.

### 4.1 Nine Proteins Differentiate Adolescents with AN/Atyp-AN from HC

We identified nine proteins suggestive of inflammation in restrictive eating disorders broadly (i.e., AN and atyp-AN) that were different from those in HC. Four proteins (LIF-R, CCL23, MMP-1, Flt3L) had higher plasma concentrations in AN/atyp-AN compared with HC, and five proteins (TRANCE, IL-18R1, HGF, CXCL1, TNFSF14) had lower plasma concentrations in AN/atyp-AN compared to HC. Of the nine proteins we identified, six (LIF-R, TRANCE, IL-18R1, HGF, CXCL1, TNFSF14) were previously identified by Nilsson et al. in an AN-only sample. Notably, the direction of the protein concentrations (*higher* LIF-R; *lower* TRANCE, IL-18R1, HGF, CXCL1, TNFSF14) are also consistent with those reported by Nilson et al. The data also show, for the first time, three additional proteins (CCL23, MMP-1, Flt3L) which show differential expression in AN and atyp-AN. CCL23 is a chemokine involved in immune cell migration and inflammation;^38^ MMP-1 is an enzyme from the matrix metalloproteinase family that degrades components of the extracellular matrix, particularly collagen;^39^ and Flt3L is a cytokine that regulates the development and function of dendritic cells, a type of immune cell.^40^ Flt3L expressed centrally is essential for maintaining brain homeostasis, and is abundantly expressed in the pyramidal cells of the hippocampus.^41^ While we only have peripheral Flt3L levels, peripheral Flt3L can influence central levels through either transport across the blood-brain barrier, indirect effects on central Flt3L, or migration via dendritic cells influenced by Flt3L.

Interestingly, all four proteins with higher plasma concentration levels in AN/atyp-AN (CCL23, LIF-R, MMP-1, Flt3L) are believed to play a pathological role in the development or progression of autoimmune rheumatoid arthritis (RA). In patients with RA, LIF-R, CCL23, MMP-1, and Flt3L expression promotes an inflammatory response and is upregulated in synovial tissue. For example, LIF-R promotes the production of pro-inflammatory cytokines (i.e., IL-6) that contribute to joint damage in RA.^42, 43^ CCL23, also known as MIP-3, is a chemokine involved in the recruitment and activation of immune cells, including monocytes, dendritic cells, and T cells, and contributes to the infiltration of inflammatory cells to the joints, promoting chronic inflammation.^44, 45^ MMP-1, an enzyme belonging to the matrix metalloproteinase family, is involved in the breakdown of extracellular matrix components, including collagen.^46, 47^

Several studies have shown bi-directional associations between AN and autoimmune^48^ disorders, including RA,^48^ and comorbid RA and AN have been reported anecdotally.^49^ However, longitudinal studies that explore the risk of RA following AN require extended longitudinal follow-up (e.g., rheumatoid average age of onset is typically much later around age 44)^48, 50, 51^ making it challenging to understand the relationship between the two diseases. Of note, the first genome-wide significant association in AN was identified in a region previously implicated in autoimmune diseases, including arthritis.^52, 53^ Genetic polymorphisms, and one’s genetic composition, play a critical role in the expression of inflammatory proteins.^54–57^ As such, the elevation of CCL23, LIF-R, MMP-1, and Flt3L observed in our study may reflect genetic contribution of previously identified genetic regions associated with AN and RA. Future studies combining genome-wide association (GWA) and plasma proteomics are necessary to determine the contribution of either genes or state-dependent traits on inflammatory biomarkers, and to further characterize the utility of these plasma biomarkers in AN diagnosis and disease progression.

### 4.2 Unique Expression Levels of IL-18R1, TRANCE, LIF-R Differentiate AN and HC

The inclusion of a group of individuals with AN *and* atyp-AN allowed us to explore proteins that may be unique to the clinical presentation of restrictive eating marked by low-weight status present in AN. We identified three proteins (*lower* IL-18R1, TRANCE; *higher* LIF-R) which were uniquely disrupted in females with AN. Our findings of lower TRANCE and higher LIF-R align with Nilsson et al., but conflict with two prior studies that reported higher concentration of TRANCE in AN than controls.^58, 59^ TRANCE is a critical regulator of bone remodeling,^60^ and the observed low levels of TRANCE among AN and within the broader AN/atyp-AN group, suggest decreased osteoclast activity, leading to reduced bone reabsorption, and altered bone remodeling,^61, 62^ that characterize clinical bone loss observed in AN.^63, 64^

Notably, individuals with atyp-AN may also be at risk for clinical bone loss, although results have been inconclusive to date,^65, 66^ likely due to heterogeneity of atyp-AN presentations.^3^ LIF-R is an inflammatory cytokine often implicated in muscle and fat wasting in cancer-related cachexia.^67^ LIF binds with a receptor that is very similar to the IL-6 receptor, both of which act via Jak2/stat3 signaling.^68, 69^ Dysregulated Jak2/stat3 signaling contributes to muscle atrophy in an estrogens dependent manner.^70, 71^ That is, only ovariectomized mice show muscle atrophy from dysreguated Jak2/stat3 signaling.^70, 71^ Estrogens deficiency is common in AN (∼60%), and may contribute to the observed inflammation patterns in the low-weight state of AN. The observed peripheral elevations of LIF-R in AN may suggest that muscle size and health cannot be maintained in this low-weight state. As we observe elevations of LIF-R among the combined AN and atyp-AN group, weight restoration above and beyond 90% of expected body weight, and restoration of menses, are likely needed for muscle restoration. LIF-R, TRANCE, and IL-18R1 are also all correlated with BMI across the entire sample; normalization of these proteins may serve as a more accurate indicator of healthy weight than BMI values in AN.

### 4.3 Twenty-Seven Proteins are Correlated with BMI Across Sample

We identified 27 proteins correlated with BMI across the entire sample. By employing the same targeted-proteomics approach (proximity extension assay) as the largest study of inflammation protein markers to date, we were able to replicate the association between inflammation protein expression levels and BMI of 18 out of 27 proteins identified previously (ADA, AXIN1, CCL11, CD5, CXCL11, CD5, CXCL1, HGF, IL10RB, IL12B, IL18, IL-18R1, IL6, LAP TGFß1, SIRT2, STAMBP, TNFRSF9, TNFS14, TRAIL, TRANCE).^32^ Importantly, the direction of the relationship was replicated across all 18 proteins.

Additionally, we identified nine novel proteins associated with BMI (*positive* association CD244, CD40, CD6, FGF-21; *negative* association CCL23, FGF-19, IL8, LIF-R, OPG). Several of these proteins, including CD40, FGF-21, FGF-19, have shown previous associations with BMI.^72–75^ CD40 is part of the TNF receptor superfamily and has been implicated in promoting inflammatory responses in various tissues and cell types.^76^ In pre-clinical rodent studies, CD40 signaling has been shown to contribute to the progression of obesity-related complications such as insulin resistance and cardiovascular disease.^77^ FGF-21 and FGF-19 are hormones secreted broadly (FGF-21) and by the small intestine (FGF-19);^78^ both play a critical role in regulating energy metabolism, glucose homeostasis, and lipid metabolism. Elevated circulating levels of FGF-21 are evident in individuals with obesity, particularly in those with obesity-related metabolic disorders such as insulin resistance, type 2 diabetes, and fatty liver disease.^79^ On the other hand, peripheral levels of FGF-19 have been shown to be lower among individuals with obesity and low FGF-19 levels have been linked to impaired glucose metabolism, insulin resistance, and dyslipidemia.^80, 81^ Interestingly, FGF-19 has been investigated as a potential therapeutic target for obesity and related metabolic disorders. Administration of exogenous FGF-19 or its analogs has shown beneficial effects on body weight, glucose homeostasis, and lipid profiles in animal studies and some clinical trials.^82, 83^

Overall, our results converge with previous studies and provide an important replication of the association between specific immune-related proteins and BMI, along with quantitative differences in these proteins in AN compared to HC. Further, we expand the previous findings through detailing these changes in AN and atyp-AN during adolescence, a critical period in the development of these conditions.

### 4.4 Strengths and Limitations

Strengths of this study include our use of a state-of-the-art multiplex proteomics approach to examine a comprehensive and validated profile of inflammation proteins; the inclusion of an adolescent sample; and inclusion of individuals with atyp-AN as well as AN. In addition, we took several steps to reduce confounding effects on blood-based biomarkers, including collection of fasting samples in the morning, placing samples immediately on ice to prevent degradation of markers, and immediate centrifugation, which has been shown to impact protein expression levels.^84, 85^ These steps allowed for more accurate and reliable measurements across participants. Our deeply phenotyped samples allowed us to conduct several complementary sensitivity analyses to ensure that our results were not driven by confounding factors such as smoking status, acute sickness, psychiatric severity, depression/anxiety, antidepressant use, or antihistamine use. These secondary analyses did not substantially change our findings, indicating that our results are robust.

Nonetheless, this study has several notable limitations. First, because the reported associations are cross-sectional, we were unable to determine if the observed alterations in protein expression are a consequence or part of the etiology of AN/atyp-AN. Given the results demonstrating protein expression levels that were unique to the low-weight AN group, the strong associations of certain protein levels with BMI, and previous findings by Nilsson and colleagues, we hypothesize that several of these inflammation findings are state-specific. Longitudinal follow-up studies will ultimately be necessary to better confirm this hypothesis. Finally, whole blood is a complex tissue and protein expression could differ based on blood processing or pre-analytic pipelines. The current study also was not designed to test cell-specific protein expression. While the inclusion of atyp-AN allowed us to determine proteins driven by a low-weight state, our sample of atyp-AN only included individuals who were at or below 90% of expected body weight. Future studies including individuals across the entire weight spectrum would improve our understanding of dysregulated inflammation in atyp-AN. In order to understand the complete picture of atyp-AN, including individuals who have restrictive eating and are at a higher weight, large sample sizes are needed. This is particularly important given the heterogeneous presentation of atyp-AN in the current literature.^3, 86^

### Conclusions

The current study represents the first to examine inflammation protein expression in adolescents with AN and atyp-AN using a methodologically rigorous approach. Our results include findings that replicate those previously published in adults with AN showing an aberrant inflammatory profile, and furthermore, expand upon those findings as we identify additional novel proteins that were not previously reported. Across both individuals with atyp-AN and AN (all with expected body weight ≤ 90%), we continued to observe disruptions in several inflammation proteins that are key for bone and muscle regulation, providing additional support for the severity of atyp-AN.

## Author Contributions

LB designed the study. KTE, EAL, and MM collected the samples. LB and CJ analyzed the data. LB, KTE, EAL, and MM wrote the manuscript, which was revised and approved by all authors.

## Acknowledgements

Data from this study was supported by R01MH103402, R03MH126143, and a Harvard Medical School Livingston Fellowship. LB is supported by NIMH (R03MH126143; K23MH127465), the Brain Behavior Research Foundation, and the International OCD Foundation. CMB is supported by NIMH (R56MH129437; R01MH120170; R01MH124871; R01MH119084; R01MH118278; R01 MH124871), Swedish Research Council (Vetenskapsrådet, award: 538-2013-8864), and the Lundbeck Foundation (Grant no. R276-2018-4581). MM is supported by NIMH (R01MH116205) and NIDDK (R01DK122581l; R01DK103946; R01DK124223). LH is supported by NIA (R01AG057505; U54AG062322), NIMH (U54MH118919; R03MH126143; R01MH128246), and the Foundation for Prader-Willi Research. KW is supported by NIMH (R01RH127259), the International OCD Foundation, Octapharma, the Fidelity Bioscience Research Initiative, and the University of California San Francisco (UCSF). EAL is supported by K24MH120568 and P30DK040561.

