## Supplemental Tables and Methods for "Identification of State Markers in Anorexia Nervosa: Replication and Extension of Inflammation Associated Biomarkers Using Multiplex Profiling in Anorexia Nervosa and Atypical Anorexia Nervosa" (from another version of this preprint: version 1, 2023-07-03)

**Supplemental Online Content**

**eMethods. Supplemental Methods**

**eFigure 1. Quantile–Quantile plots for Normality of Protein Expression Levels**

### eTable 1. Additional Clinical Characteristics of the Sample.

### eTable 2. Proteins included in analysis.

### eFigure 2. PCA summary plot on the 73 proteins to identify outliers.

### eMethods Supplement

Participant Criteria and Study Procedures

1. Parent Study Procedures

1.1. Parent Studies Overview

Data in the current manuscript were obtained from two parent studies:

(1) A longitudinal cohort study on the neurobiology of low-weight eating disorders (LWED) (R01MH103402). Females with anorexia nervosa (AN), atypical AN (atyp-AN), avoidant restrictive food intake disorder (ARFID), and healthy controls with no lifetime history of eating disorders (HC) participated in this 18-month prospective study aimed at exploring homeostatic and hedonic food motivation pathways as predictors of eating disorder trajectories, including hormones, neural circuitry, and psychological symptoms.

(2) A longitudinal cohort study on the neurobiology of ARFID (R01 MH108595). Youth (male and female) with ARFID and HC with no lifetime history of eating disorders participated in this two-year study aimed at exploring underlying neural, cognitive, neuroendocrine, physiological and behavioral mechanisms of ARFID.

1.2. Parent Study Hypotheses

*Parent Study 1*

Hypothesis 1: Altered food motivation pathways (ie, hormones, neural circuits, and psychological symptoms) underly patterns of restriction, binge eating, and purging, and differentiates adolescents with low-weight eating disorders from healthy, normal-weight controls.

Hypothesis 2: Within adolescents with low weight eating disorders followed longitudinally, 18-month outcome (ie, dietary restriction, binge/purge frequency, and weight changes) is determined by food motivation pathways (ie, hormones, neural circuits, and psychological symptoms) at baseline and changes over time.

*Parent Study 2*

Hypothesis 1: Avoidant/restrictive eating pathology is associated with increased

fear/threat response, impaired homeostatic appetite, and impaired sensory processing/perception.

Hypothesis 2: Within youth with ARFID followed longitudinally, two-year outcomes (e.g., avoidant/restrictive eating pathology) is associated with RDoC constructs of acute threat/fear, homeostatic appetite, and perception/sensory processing at baseline.

1.3. Participant Selection Criteria for current study

*Females with low-weight eating disorders (LWED)*

Inclusion criteria

1. Female aged 10 to 22 years old

2. Eating disorder characterized by (i) low body weight, which is defined as less than 90% of median BMI for age and sex, or less than 90% of ideal body weight for height (IBW, defined as the weight corresponding to the participant’s height percentile for age/bone age), and (ii) restrictive eating, binge eating more than once a month, purging more than once a month, excessive exercise more than once a month, and/or in treatment for an eating disorder, to be evaluated during the initial clinical assessment.

(i) Height and weight were collected to calculate BMI, percent median BMI, and percent predicted body weight for age and height. The Centers for Disease Control and Prevention (CDC) growth charts were used to calculate the percentage of median BMI and percentage of predicted body weight, where 20 years was used as the reference age for any participant older than 20 years (as the CDC charts only go to 20 years).

(ii)The LIFE-EAT-3 was used to evaluate the course of eating disorder pathology and psychosocial functioning on a weekly basis across time. According to a scale from "1" to "6," where "1" and "2" denote no/minimal symptoms, "3" or "4" denote moderate symptoms, and "5" or "6" denote severe symptoms, the psychiatric status rating (PSR), the main measurement, evaluated the course of eating disorder symptoms on a weekly basis. The American Psychiatric Association Level of Care Guidelines for Patients with Eating Disorders were used to organize the subjects' self-reported treatment status by level of care, which included outpatient, intensive outpatient, partial hospitalization/full-day program, and residential treatment. Notably, we allowed outpatient treatment to be defined by the subject and included a variety of therapy modalities (including CBT-E, FBT, hypnotherapy, psychodynamic therapy, group therapy, vital/weight checks by a medical provider), delivered by a variety of providers (such as Ph.D. level psychologists, masters level therapists, dietitians, psychiatrists, school counselors), with varying frequencies (such as once per week, bi-weekly, monthly).

Individuals with an eating disorder were diagnosed with either AN or atyp-AN.

2.1. AN diagnostic criteria by EDE/LIFE-EAT-3:

- BMI percentile of ≤ 10 for adolescents or BMI was ≤ 18.5 for adults
- LIFE-EAT PSR score ≥ 3 (indicating moderate symptoms) for the previous three months

2.2. Atyp-AN diagnostic criteria by EDE/LIFE-EAT-3:

- AN criterion by EDE/LIFE-EAT-3
- BMI percentile > 10 if < 18 years old, or had BMI > 18.5 if ≥ 18 years old

Exclusion criteria

1. Use of systemic hormones, pregnancy or breastfeeding within eight weeks

2. Use of Depo-Provera within three months

3. History of psychosis by Schedule for Affective Disorders and Schizophrenia for School Age Children-Present and Lifetime Version (KSADS-PL)

4. Substance or alcohol use disorder active within the past month by KSADS-PL 5. Diabetes mellitus

6. Hematocrit (Hct) < 30.0%

7. Potassium (K) < 3.0 mmol/L

8. Gastrointestinal tract surgery (including gastrectomy, gastric bypass surgery, and small or large bowel resection)

9. Other medical explanation for low weight (e.g., brain tumor)

10. Active suicidal ideation

*Healthy control females*

Inclusion criteria

1. Female aged 10 to 22 years old

2. BMI percentile for age and sex between 25th and 85th percentile

3. Matched to eating disorder cohort to be within 1 Tanner stage

4. Having regular menses for those more than 2 years from menarche

5. No pubertal delay (pubertal delay includes menarche at > 16 years, thelarche at > 13 years).

Exclusion criteria

1. Use of systemic hormones, pregnancy, or breastfeeding within eight weeks of screening

2. Use of Depo-Provera within three months of screening

3. History of lifetime psychiatric diagnoses by KSADS-PL

4. Diabetes mellitus

5. Hematocrit (Hct) < 30.0%

6. Potassium (K) < 3.0 mmol/L

7. Gastrointestinal tract surgery (including gastrectomy, gastric bypass surgery, and small or large bowel resection)

8. Active suicidal ideation

9. Excessive exercise within the last 3 months (having run > 25 miles in any one week or having exercised >10 hours in any one week)

10. Vegetarianism

11. Familial history of AN or atyp-AN in first-degree relatives

1.4. Study Procedures for Parent Study 1 and 2

*Individuals presented to morning visits in their fasting states*

Pre-meal procedures overview

1. History and physical exam with evaluation of height, weight, and Tanner stage

2. Assessments of dietary habits and physical activity levels with 4-day food record and Paffenbarger Questionnaire

3. Blood draw for fasting hormones

4. MRI pre-meal

5. Pre-meal, self-reported psychological symptoms

Study meal

Subjects consumed a standardized mixed meal with a macronutrient composition of approximately 20% fat, 60% carbohydrates, and 20% protein, totaling approximately 400-kilocalories.

Post-meal procedure overview

1. Serial blood draws for postprandial hormones

2. Magnetic Resonance Imaging 30 minutes post-meal

3. Psychological symptom reports post-meal

Behavioral measures and clinical interviews used in screening and/or main study visits:

*Parent Study 1 and 2*

1. Schedule for Affective Disorders and Schizophrenia for School Age Children-Present and Lifetime Version (K-SADS-PL)
2. Annett Hand Preference Questionnaire
3. Appetite Visual Analog Scales (VAS)
4. Saliency VAS
5. Clinical Impairment Assessment (CIA)
6. Eating Disorder Examination Questionnaire (EDE-Q)
7. Eating Disorders Longitudinal Interval Follow-up Evaluation 3 (LIFE-EAT 3)
8. Beck Depression inventory II (BDI-II) (or Children’s Depression Inventory, CDI-2)
9. Beck Anxiety Inventory (or BAI–Youth, BAI-Y)
10. State Trait Anxiety Inventory (Trait only) (or STAI for Children, Trait only, STAIC)
11. Eating Inventory (EI)
12. Adolescent/Adult Sensory Profile (AASP)
13. Strengths and Difficulties Questionnaire (SDQ)
14. Pica ARFID Rumination Disorder Interview (PARDI)
15. Food Neophobia Questionnaire (FNQ)
16. Adult Eating Behaviour Questionnaire (AEBQ)
17. ARFID prototype matching form
18. Toronto Alexithymia Scale (TAS-20)
19. Temporal Experience of Pleasure Scale (TEPS)
20. Behavioral Inhibition and Activation Scales (BIS/BAS)
21. Liebowitz Social Anxiety Scale (LSAS-SR)
22. Power of Food Scale (PFS)
23. Regional Taste Intensity Test
24. Paffenbarger Activity Assessment
25. 24-hour food recall
26. Four-Day Food Record
27. Delay Discounting Task

*Parent Study 1 Only*

1. Eating Expectancy Inventory (EEI)
2. Eating Disorder Inventory-3 (EDI-3)
3. Child Development Inventory
4. Alcohol Use Disorders Identification Test (AUDIT)
5. Drug Use Questionnaire (DAST-10)
6. Cookie Taste Test
7. Eating Disorder Examination (EDE)
8. Implicit Association Test (IAT)
9. Go/No-Go (GNG) Task

*Parent Study 2 Only*

1. Eating Disorders Assessment for DSM-5 (EDA-5)
2. Eating Disorder Inventory-3 (EDI-3, Drive for Thinness subscale only)

Optional procedures with additional consenting required

1. Saliva or blood collection for DNA

2. Hair collection to assess cortisol concentration

3. Stool collection for microbiome profiling

4. Dual-energy X-ray absorptiometry (DXA) for bone mass density and body composition

### eFigure 1. Quantile–Quantile plots for Normality of Protein Expression Levels

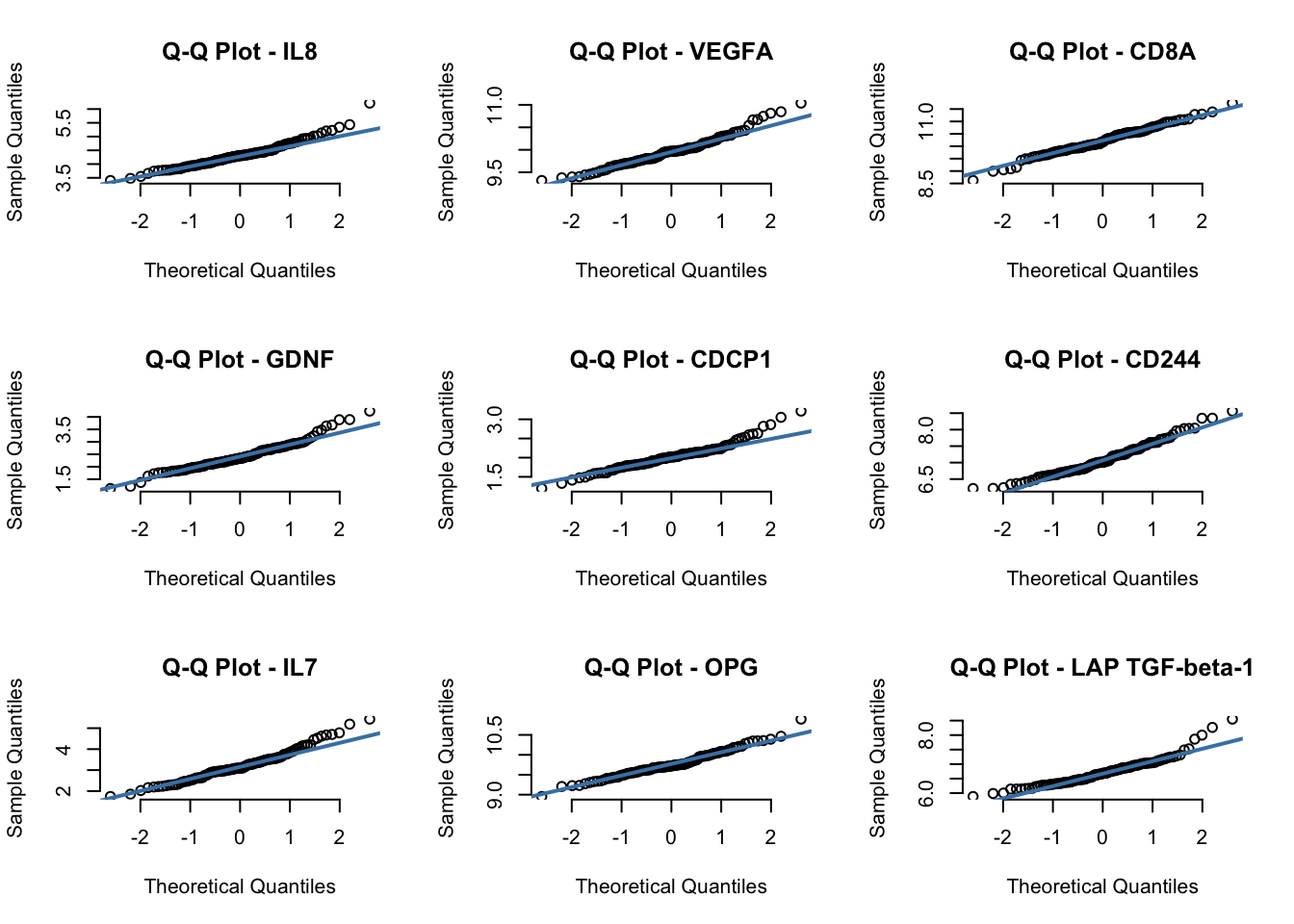

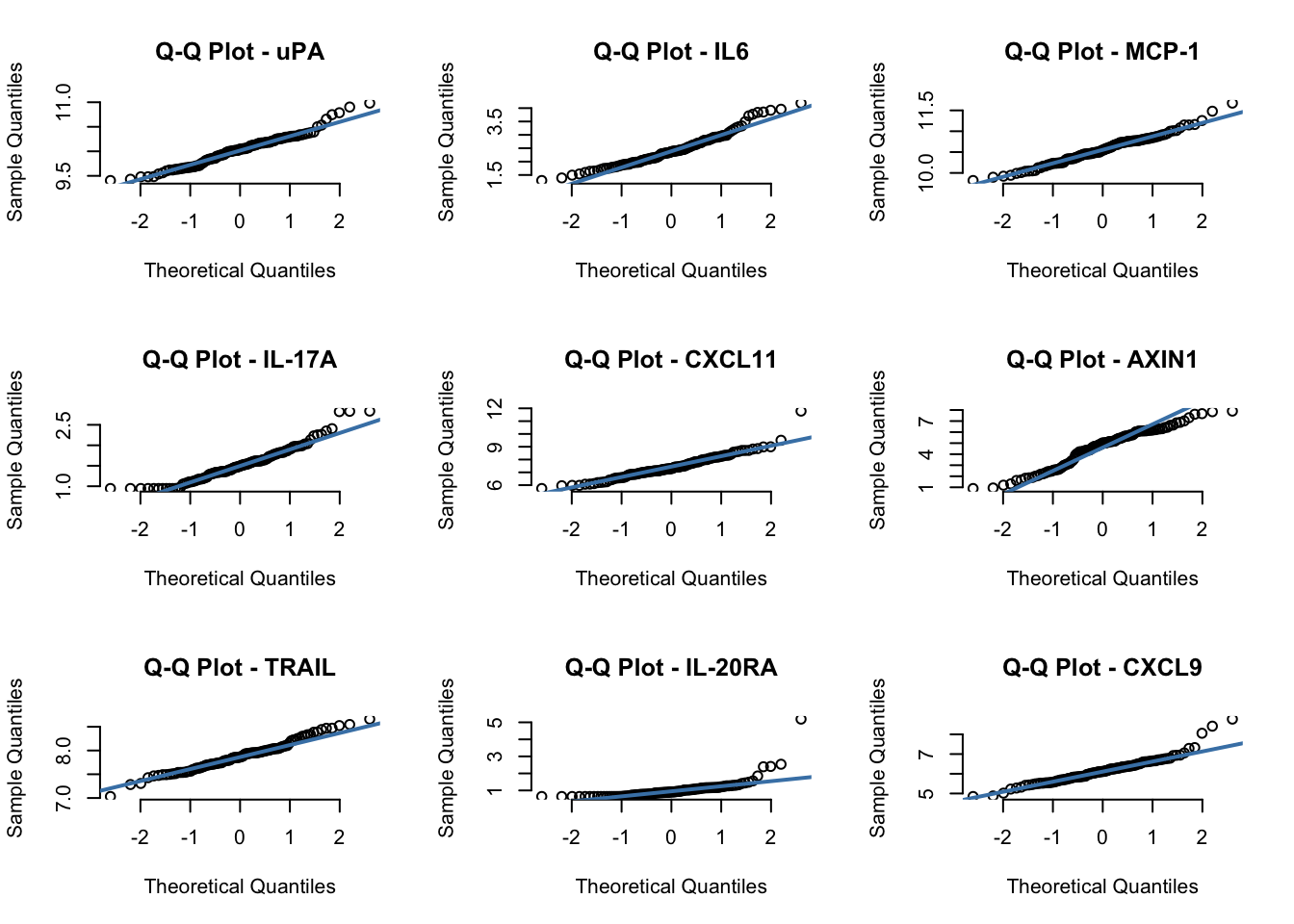

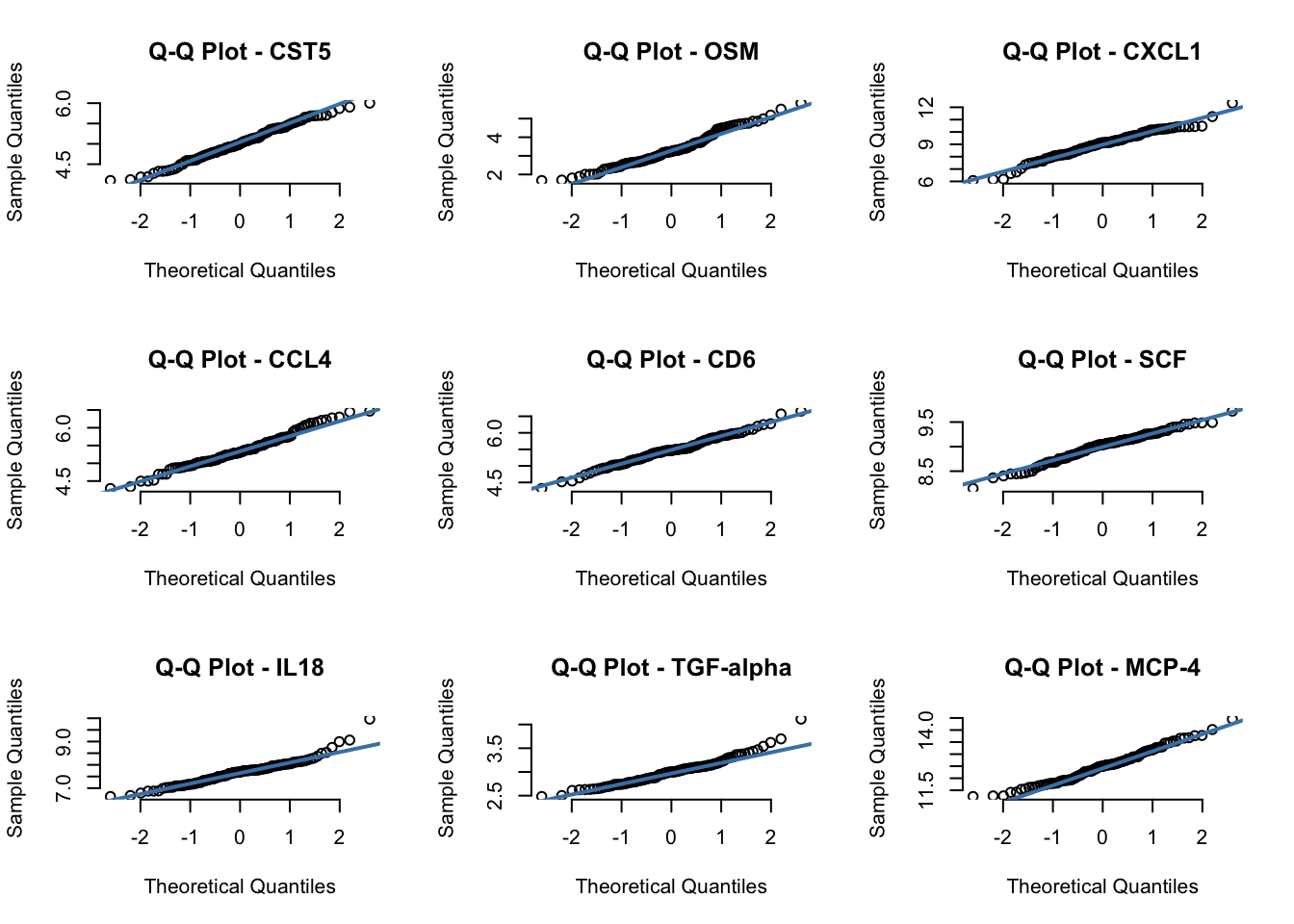

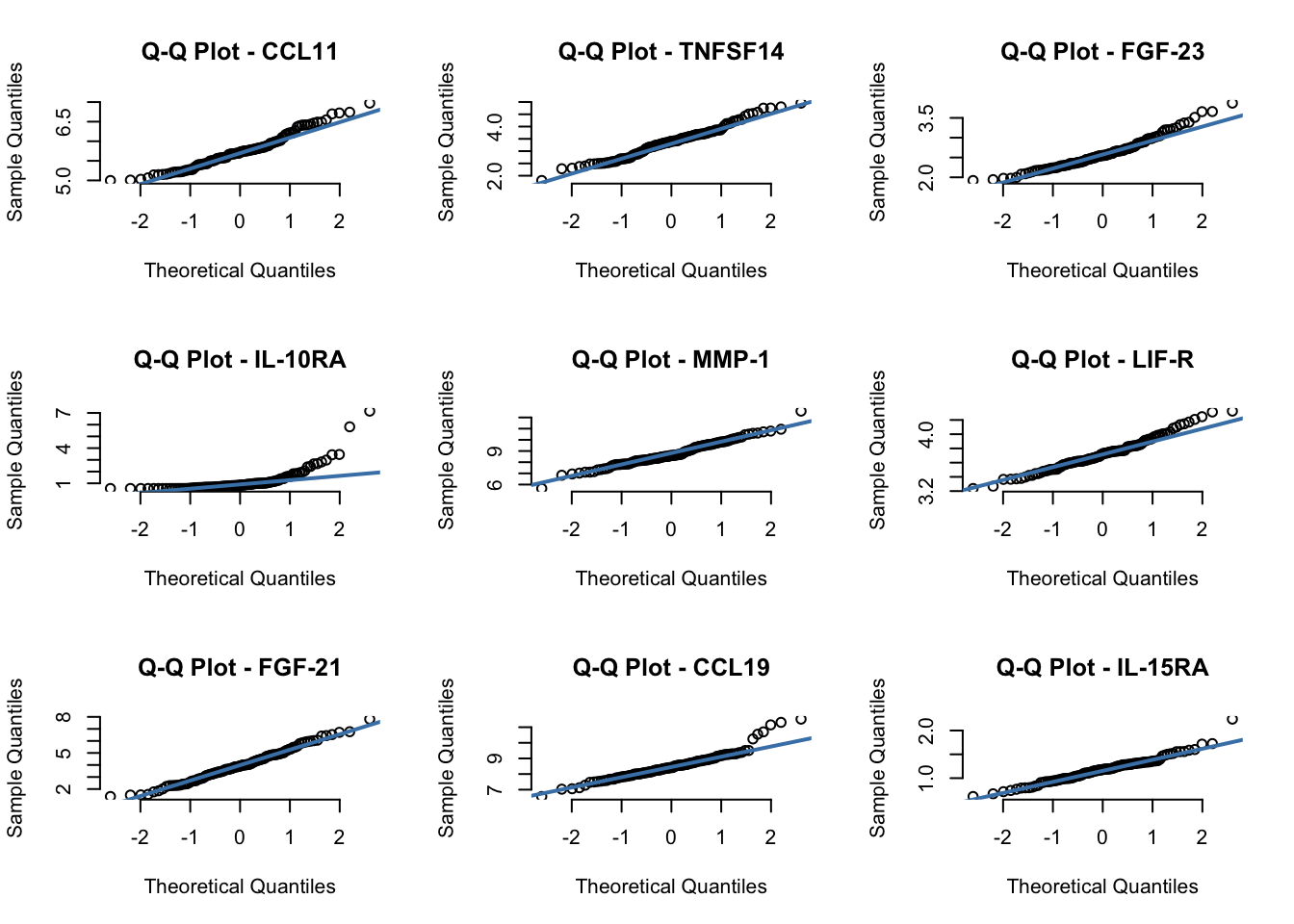

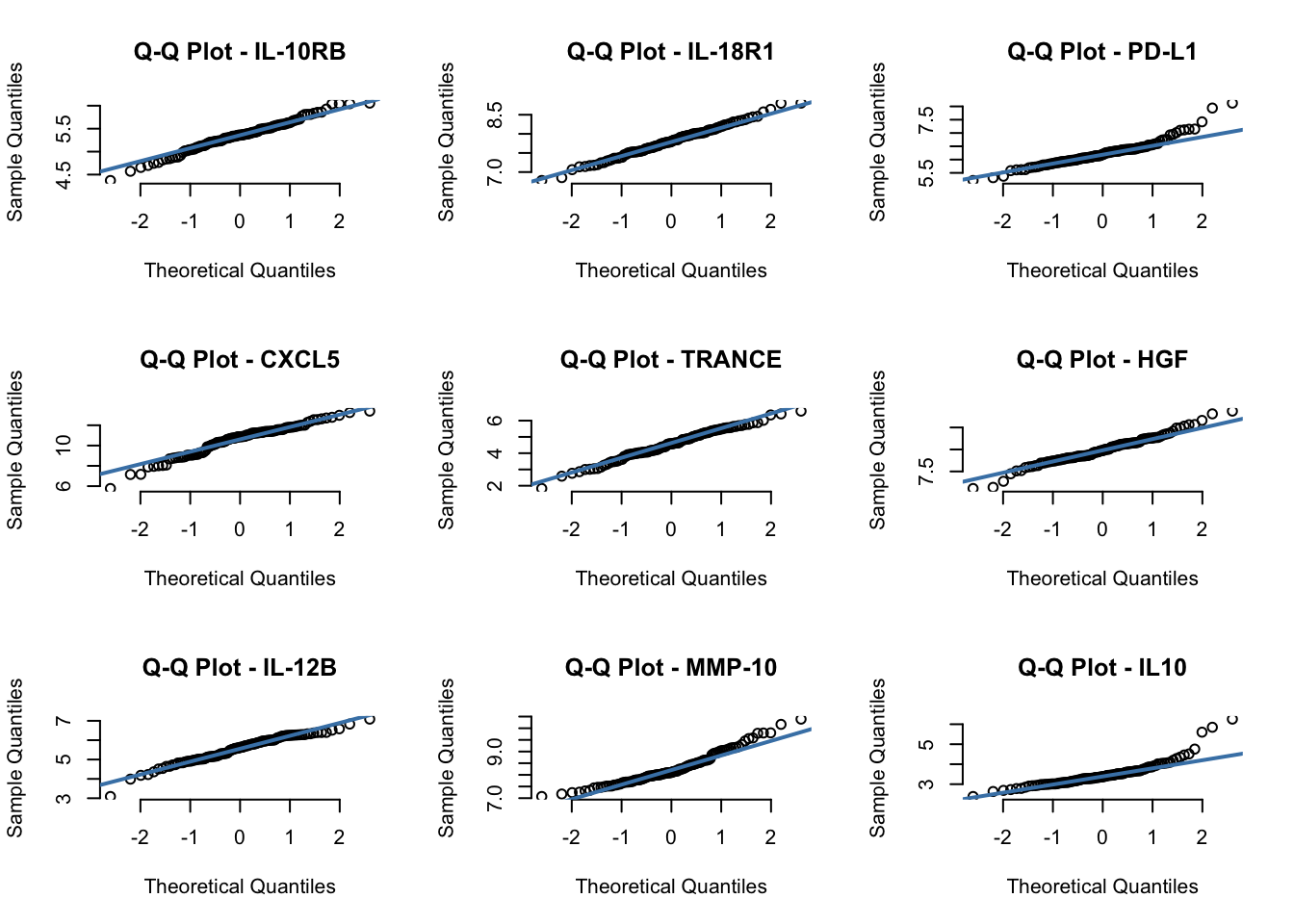

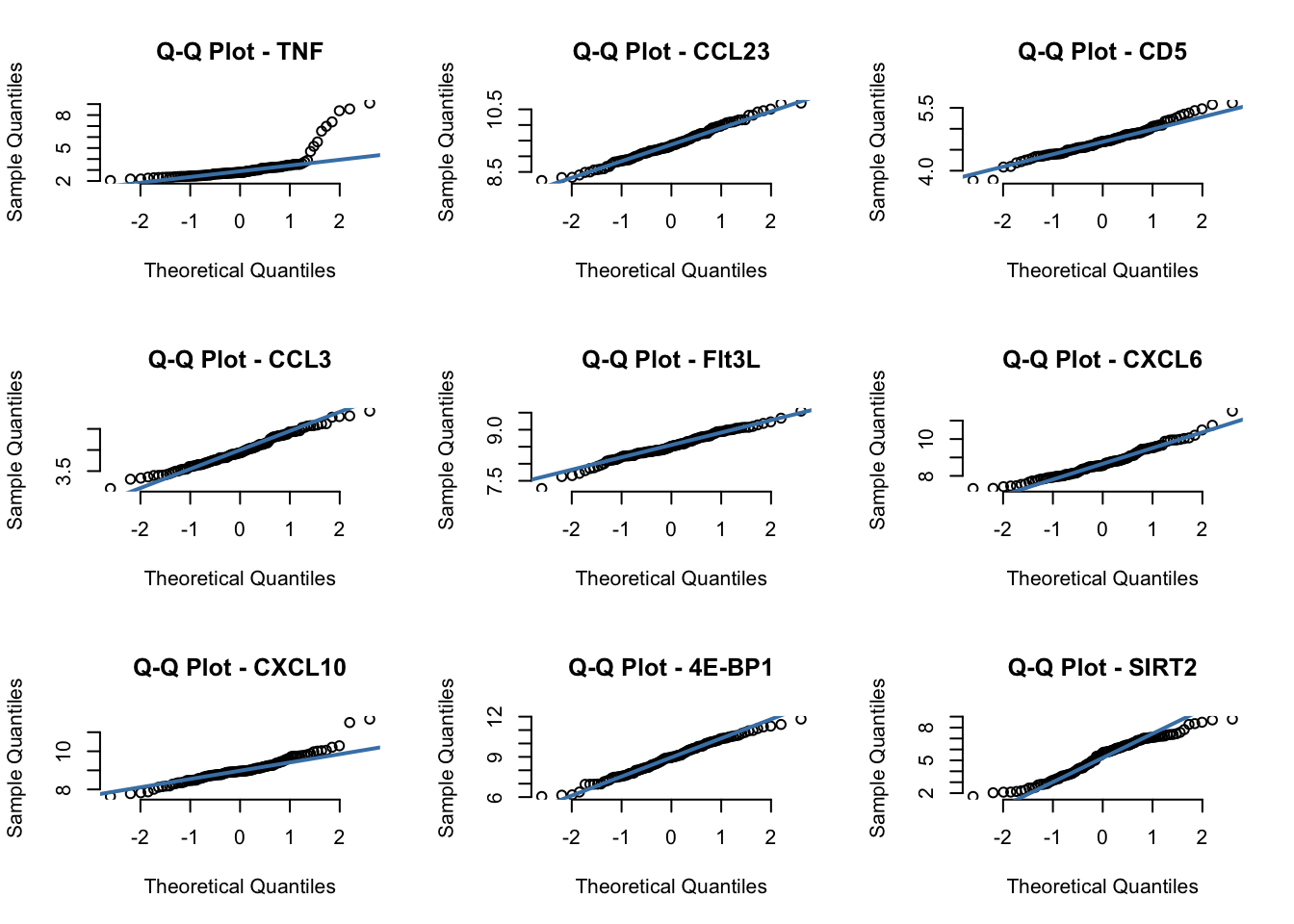

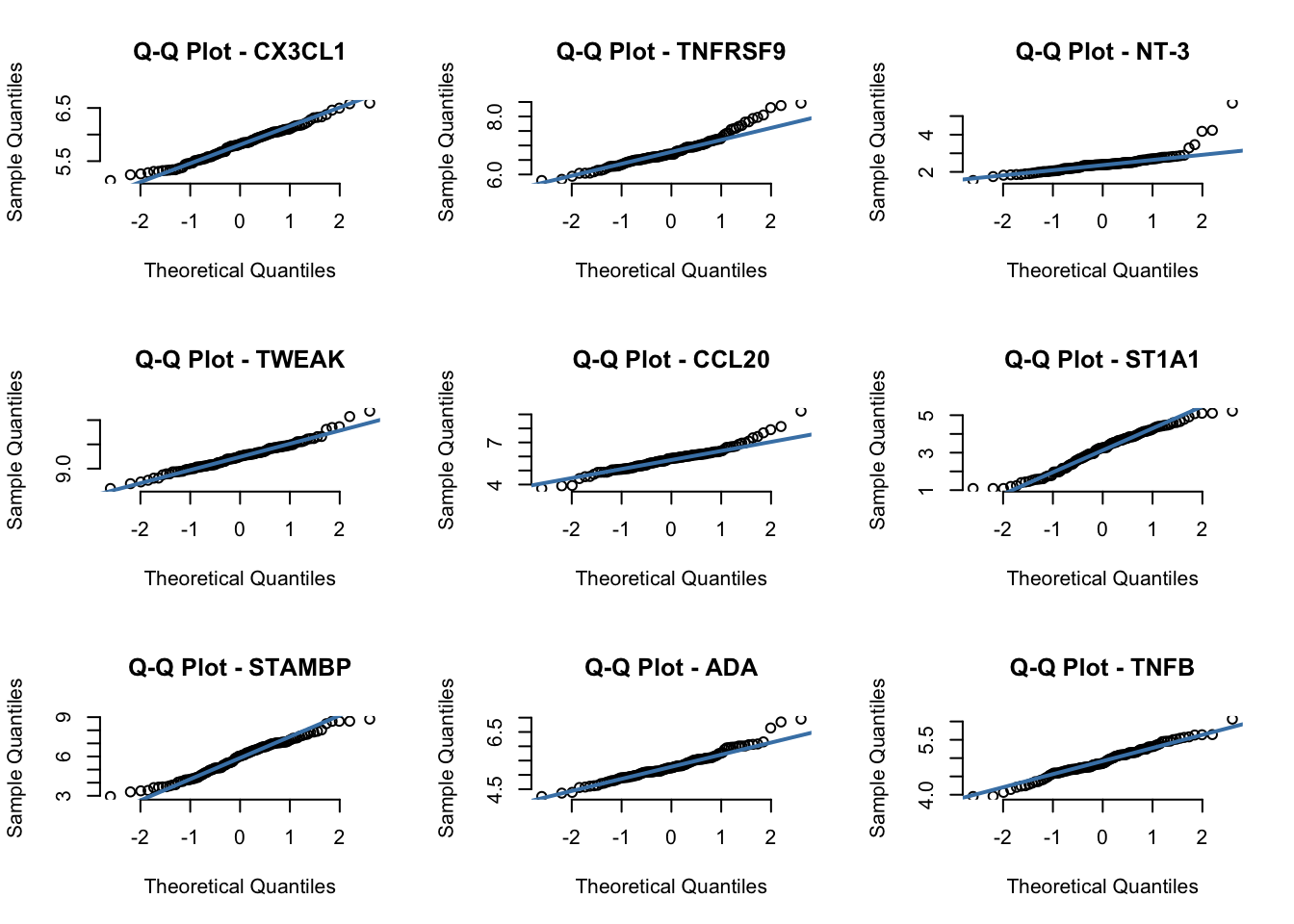

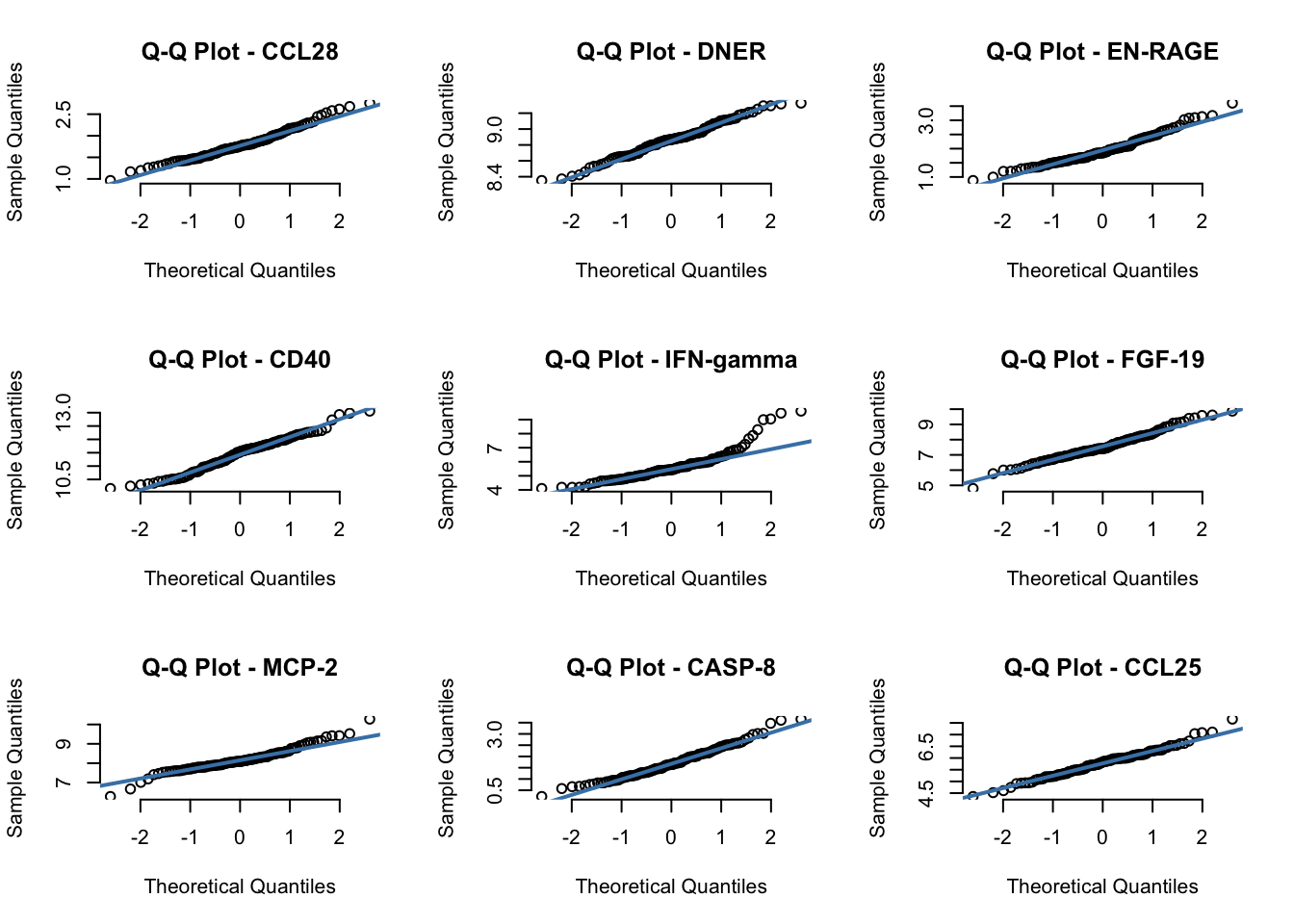

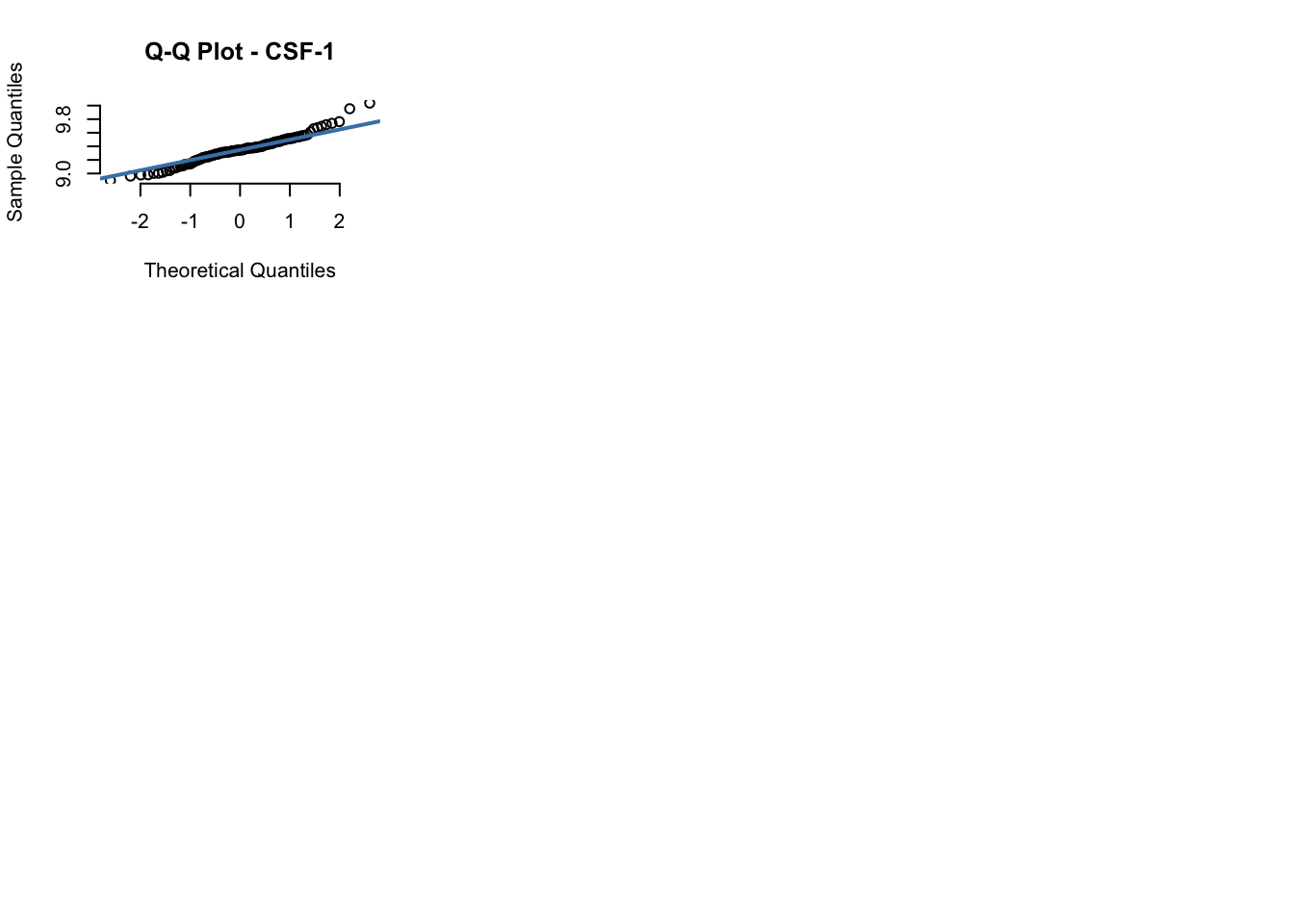

### eTable 1. Additional Clinical Characteristics of the Sample.

|  | **AN**  **(N=38)** | **atyp-AN**  **(N=26)** | **HC**  **(N=44)** | **Overall**  **(N=108)** |
| --- | --- | --- | --- | --- |
| **Body Temperature (°F)** |  |  |  |  |
| Mean (SD) | 97.9 (0.634) | 97.9 (0.732) | 98.1 (0.724) | 98.0 (0.696) |
| **Anxiety t-score** |  |  |  |  |
| Mean (SD) | 67.7 (12.8) | 66.5 (11.9) | 44.7 (8.09) | 58.1 (15.5) |
| **Depression t-score (** |  |  |  |  |
| Mean (SD) | 59.2 (7.72) | 62.8 (9.25) | 43.1 (7.01) | 53.4 (11.7) |
| Missing | 0 (0%) | 1 (3.8%) | 0 (0%) | 1 (0.9%) |
| **Mood Disorders** |  |  |  |  |
| Yes | 5 (13.2%) | 6 (23.1%) | 0 (0.0%) | 12 (11.1%) |
| No | 33 (86.8%) | 20 (76.9%) | 44 (100.0%) | 96 (88.9%) |
| **Anxiety Disorders** |  |  |  |  |
| Yes | 21 (55.3%) | 15 (57.7%) | 0 (0%) | 36 (33.3%) |
| No | 17 (44.7%) | 11 (42.3%) | 44 (100%) | 72 (66.7%) |
| **PTSD** |  |  |  |  |
| Yes | 2 (5.3%) | 0 (0%) | 0 (0%) | 2 (1.9%) |
| No | 36 (94.7%) | 26 (100%) | 44 (100%) | 106 (98.1%) |
| **ADHD** |  |  |  |  |
| Yes | 1 (2.6%) | 1 (3.8%) | 0 (0%) | 2 (1.9%) |
| No | 37 (97.4%) | 25 (96.2%) | 44 (100%) | 106 (98.1%) |
| **ODD** |  |  |  |  |
| Yes | 0 (0%) | 0 (0%) | 0 (0%) | 0 (0%) |
| No | 38 (100%) | 26 (100%) | 44 (100%) | 108 (100%) |
| **ASD** |  |  |  |  |
| Yes | 0 (0%) | 0 (0%) | 0 (0%) | 0 (0%) |
| No | 38 (100%) | 26 (100%) | 44 (100%) | 108 (100%) |
| **SUD** |  |  |  |  |
| Yes | 0 (0%) | 0 (0%) | 0 (0%) | 0 (0%) |
| No | 38 (100%) | 26 (100%) | 44 (100%) | 108 (100%) |
| **AUD** |  |  |  |  |
| Yes | 1 (2.6%) | 0 (0%) | 0 (0%) | 1 (0.9%) |
| No | 37 (97.4%) | 26 (100%) | 44 (100%) | 107 (99.1%) |
| **Antidepressant Medications** |  |  |  |  |
| Yes | 19 (50.0%) | 9 (34.6%) | 0 (0%) | 28 (25.9%) |
| No | 19 (50.0%) | 17 (65.4%) | 44 (100%) | 80 (74.1%) |
| **Anti-anxiety Medications** |  |  |  |  |
| Yes | 6 (15.8%) | 10 (38.5%) | 0 (0%) | 16 (14.8%) |
| No | 32 (84.2%) | 16 (61.5%) | 44 (100%) | 92 (85.2%) |
| **Stimulants** |  |  |  |  |
| Yes | 3 (7.9%) | 0 (0%) | 0 (0%) | 3 (2.8%) |
| No | 35 (92.1%) | 26 (100%) | 44 (100%) | 105 (97.2%) |
| **Mood Stabilizers** |  |  |  |  |
| Yes | 1 (2.6%) | 1 (3.8%) | 0 (0%) | 2 (1.9%) |
| No | 37 (97.4%) | 25 (96.2%) | 44 (100%) | 106 (98.1%) |
| **Anti Obsessive Agents** |  |  |  |  |
| Yes | 1 (2.6%) | 0 (0%) | 0 (0%) | 1 (0.9%) |
| No | 37 (97.4%) | 26 (100%) | 44 (100%) | 107 (99.1%) |
| **Antipsychotics** |  |  |  |  |
| Yes | 3 (7.9%) | 2 (7.7%) | 0 (0%) | 5 (4.6%) |
| No | 35 (92.1%) | 24 (92.3%) | 44 (100%) | 103 (95.4%) |
| **Other Medications** |  |  |  |  |
| Yes | 0 (0%) | 1 (3.8%) | 0 (0%) | 1 (0.9%) |
| No | 38 (100%) | 25 (96.2%) | 44 (100%) | 107 (99.1%) |

AN = anorexia nervosa, atyp-AN = atypical anorexia nervosa, HC = healthy control, SD = standard deviation, PTSD = post-traumatic stress disorder, ADHD = attention-deficit/hyperactivity disorder, ODD = oppositional defiant disorder, ASD = autism spectrum disorder, SUD = substance use disorder, AUD = alcohol use disorder. No group differences emerged between AN and atyp-AN for the frequency of clinical characteristics reported above. As expected, individuals with AN/atyp-AN reported significantly higher levels of anxiety/depression symptom, and had significantly higher frequencies of mood disorders, anxiety disorders, antidepressant use, and anti-anxiety use than HC. Anxiety symptoms were measured by the State Trait Anxiety Inventory (Trait only) (or STAI for Children, Trait only, STAIC). Depression Symptoms were measured by the Beck Depression inventory II (BDI-II) (or Children’s Depression Inventory, CDI-2).

### eTable 2. Proteins included in analysis.

| **Protein** | **Protein full name** | **UniProt** | **OlinkID** |
| --- | --- | --- | --- |
| 4E-BP1 | Eukaryotic translation initiation factor  4E-binding protein 1 | Q13541 | OID00536 |
| ADA | Adenosine Deaminase | P00813 | OID00560 |
| AXIN1 | Axin-1 | O15169 | OID00487 |
| CASP-8 | Caspase-8 | Q14790 | OID00550 |
| CCL11 | Eotaxin | P51671 | OID00505 |
| CCL19 | C-C motif chemokine 19 | Q99731 | OID00513 |
| CCL20 | C-C motif chemokine 20 | P78556 | OID00556 |
| CCL23 | C-C motif chemokine 23 | P55773 | OID00530 |
| CCL25 | C-C motif chemokine 25 | O15444 | OID00551 |
| CCL28 | C-C motif chemokine 28 | Q9NRJ3 | OID00539 |
| CCL3 | C-C motif chemokine 3 | P10147 | OID00532 |
| CCL4 | C-C motif chemokine 4 | P13236 | OID00498 |
| CD244 | Natural killer cell receptor 2B4 | Q9BZW8 | OID00477 |
| CD40 | CD40L receptor | P25942 | OID00542 |
| CD5 | T-cell surface glycoprotein CD5 | P06127 | OID00531 |
| CD6 | T cell surface glycoprotein CD6 isoform | P30203 | OID00499 |
| CD8A | T-cell surface glycoprotein CD8 alpha chain | P01732 | OID05124 |
| CDCP1 | CUB domain-containing protein 1 | Q9H5V8 | OID00476 |
| CSF-1 | Macrophage colony-stimulating factor 1 | P09603 | OID00562 |
| CST5 | Cystatin D | P28325 | OID00491 |
| CX3CL1 | Fractalkine | P78423 | OID00552 |
| CXCL1 | C-X-C motif chemokine 1 | P09341 | OID00496 |
| CXCL10 | C-X-C motif chemokine 10 | P02778 | OID00535 |
| CXCL11 | C-X-C motif chemokine 11 | O14625 | OID00486 |
| CXCL5 | C-X-C motif chemokine 5 | P42830 | OID00520 |
| CXCL6 | C-X-C motif chemokine 6 | P80162 | OID00534 |
| CXCL9 | C-X-C motif chemokine 9 | Q07325 | OID00490 |
| DNER | Delta and Notch-like epidermal growth factor-related receptor | Q8NFT8 | OID01213 |
| EN-RAGE | Protein S100-A12 | P80511 | OID00541 |
| FGF-19 | Fibroblast growth factor 19 | O95750 | OID00545 |
| FGF-21 | Fibroblast growth factor 21 | Q9NSA1 | OID00512 |
| FGF-23 | Fibroblast growth factor 23 | Q9GZV9 | OID00507 |
| Flt3L | Fms-related tyrosine kinase 3 ligand | P49771 | OID00533 |
| GDNF | Glial cell line-derived neurotrophic factor | P39905 | OID00475 |
| HGF | Hepatocyte growth factor | P14210 | OID00522 |
| IFN-gamma | Interferon gamma | P01579 | OID05547 |
| IL-10RA | Interleukin-10 receptor subunit alpha | Q13651 | OID00508 |
| IL-10RB | Interleukin-10 receptor subunit beta | Q08334 | OID00515 |
| IL-12B | Interleukin-12 subunit beta | P29460 | OID00523 |
| IL-15RA | Interleukin-15 receptor subunit alpha | Q13261 | OID00514 |
| IL-17A | Interleukin-17A | Q16552 | OID00485 |
| IL-18R1 | Interleukin-18 receptor 1 | Q13478 | OID00517 |
| IL-20RA | Interleukin-20 receptor subunit alpha | Q9UHF4 | OID00489 |
| IL10 | Interleukin-10 | P22301 | OID00528 |
| IL18 | Interleukin-18 | Q14116 | OID00501 |
| IL6 | Interleukin-6 | P05231 | OID00482 |
| IL7 | Interleukin-7 | P13232 | OID00478 |
| IL8 | Interleukin-8 | P10145 | OID00471 |
| LAP TGF-beta-1 | Latency-associated peptide transforming growth factor beta-1 | P01137 | OID00480 |
| LIF-R | Leukemia inhibitory factor receptor | P42702 | OID00511 |
| MCP-1 | Monocyte chemotactic protein 1 | P13500 | OID00484 |
| MCP-2 | Monocyte chemotactic protein 2 | P80075 | OID00549 |
| MCP-4 | Monocyte chemotactic protein 4 | Q99616 | OID00504 |
| MMP-1 | Matrix metalloproteinase-1 | P03956 | OID00510 |
| MMP-10 | Matrix metalloproteinase-10 | P09238 | OID00527 |
| NT-3 | Neurotrophin-3 | P20783 | OID00554 |
| OPG | Osteoprotegerin | O00300 | OID00479 |
| OSM | Oncostatin-M | P13725 | OID00494 |
| PD-L1 | Programmed cell death 1 ligand 1 | Q9NZQ7 | OID00518 |
| SCF | Stem cell factor | P21583 | OID00500 |
| SIRT2 | SIR2-like protein 2 | Q8IXJ6 | OID00538 |
| ST1A1 | Sulfotransferase 1A1 | P50225 | OID00557 |
| STAMBP | STAM-binding protein | O95630 | OID00558 |
| TGF-alpha | Transforming growth factor alpha | P01135 | OID00503 |
| TNF | Tumor necrosis factor | P01375 | OID05548 |
| TNFB | TNF-beta | P01374 | OID00561 |
| TNFRSF9 | Tumor necrosis factor receptor superfamily member 9 | Q07011 | OID00553 |
| TNFSF14 | Tumor necrosis factor ligand superfamily member 14 | O43557 | OID00506 |
| TRAIL | TNF-related apoptosis-inducing ligand | P50591 | OID00488 |
| TRANCE | TNF-related activation-induced cytokine | O14788 | OID00521 |
| TWEAK | Tumor necrosis factor (Ligand) superfamily, member 12 | O43508 | OID00555 |
| VEGFA | Vascular endothelial growth factor A | P15692 | OID00472 |
| uPA | Urokinase-type plasminogen activator | P00749 | OID00481 |

### eFigure 2. PCA summary plot on the 73 proteins to identify outliers. s72 and s77 from the HC group were identified as outliers and removed from further analysis.

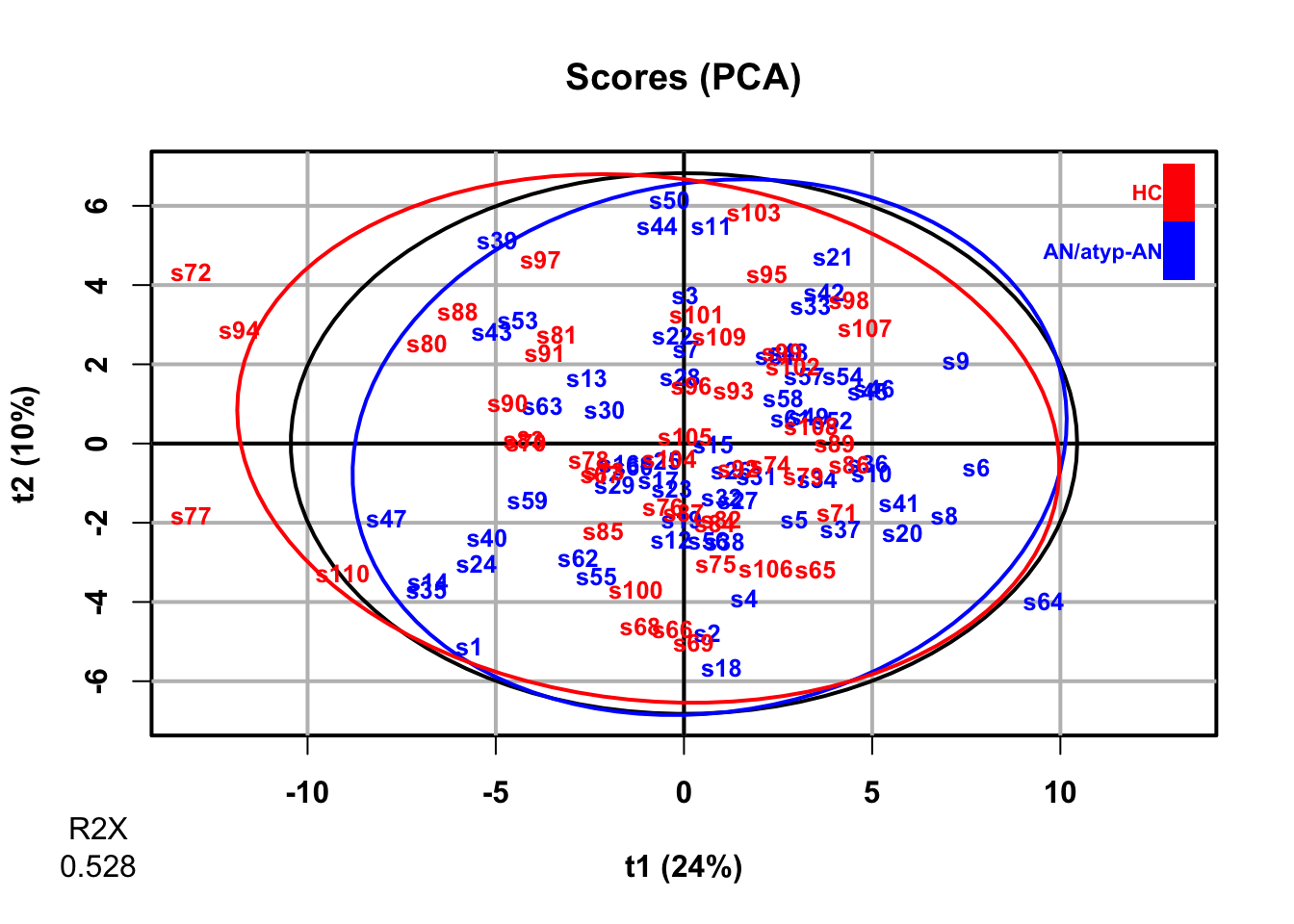
